# The mechanism of regulation of prokaryotic Na^+^ channels by their C-terminal domain

**DOI:** 10.64898/2026.11.20.768188

**Authors:** Angel Zavala-Custodio, Carlos Gonzalez, Vikram Khipple Mulligan, S. Mahsa Mofidi, Ashley Vazhavilla, Gerhard Hummer, Sonya M. Hanson, Andrés Jara-Oseguera

## Abstract

Prokaryotic voltage-gated Na^+^-channels (BacNa_V_s) closely resemble neuronal Na_V_ channels in their structure and gating behavior and are found in multiple bacterial lineages, yet little is known about their biological function. In Na_V_Sp1 from the marine bacterium *Ruegeria pomeroyi*, the cytoplasmic C-terminal domain (CTD) was proposed to function as a temperature sensor. Here, we use electrophysiology and computational methods to establish how the CTD can tune the temperature- and voltage-dependence of the Na_V_Sp1 channel. We show that although the CTD of Na_V_Sp1 is not a temperature sensor, its conformational dynamics are directly coupled to the gating of the pore. Additionally, we find that interactions between the CTD and the transmembrane domain can reduce or completely suppress the response of Na_V_Sp1 to temperature by stabilizing the open state of the channel. Although BacNa_V_ channels share low sequence identity in their CTDs, we provide evidence suggesting that most are structurally similar to Na_V_Sp1 through the mining and modeling of more than 800 sequences from distinct bacteria. The mechanism we described could therefore be important for BacNa_V_ channel regulation in response to environmental factors like temperature in a wide number of bacterial species.

**Significance:** Bacteria from soil, aquatic, plant, and animal environments have genes coding for voltage-gated Na^+^ (BacNa_V_) channels. Understanding how environmental stimuli regulate BacNa_V_s is crucial to elucidate their function in bacterial physiology, with implications for global climate and industrial bioproduction; for example, marine bacteria like *Ruegeria pomeroyi* contribute to cloud formation by producing dimethyl sulfide, a gas that nucleates water droplets in the atmosphere. Here we describe how the C-terminal domain of Na_V_Sp1 from *Ruegeria pomeroyi* regulates its channel response to voltage and temperature, and we provide evidence suggesting that this mechanism is conserved in major fraction of BacNa_V_s. Our findings and structural modeling of over 800 BacNa_V_ sequences provide a valuable framework to investigate the function of BacNa_V_s in prokaryotes.

## Introduction

Activation and inactivation of voltage-gated Na^+^ (Na_V_) channels in neurons drives the depolarizing phase of action potentials. The voltage-dependent activity of human Na_V_ channels is subject to tight modulation through association with auxiliary β subunits and a multitude of cellular mediators (1). Although voltage-gated channels are recognized for their role in neuronal excitability, these proteins have existed since before prokaryotes and eukaryotes diverged over a billion years ago. Bacterial Na_V_ channels (BacNa_V_s) possess a similar transmembrane domain fold and voltage-sensing mechanism, and give rise to currents that activate and inactivate with comparable kinetics and voltage-dependence as the neuronal channels (2–5). In alkaliphilic bacteria that populate environments that are poor in H^+^ but have high Na^+^ ion concentrations, BacNa_V_ channels were shown to have a role in motility, chemotaxis, and pH regulation (6)(7), utilizing Na^+^ instead of H^+^ to power these cellular processes. However, little else is known about the function or regulation of these channels in other species of bacteria (8). Taxonomic classification of 825 non-redundant BacNa_V_ channel sequences identifies them in at least three major classes of bacteria **(Fig. 1a, Dataset S01)** that include species populating diverse environments (2) and having distinct lifestyles, metabolisms, and physiologies **(Fig. 1b, Dataset S02)**. Some of these species are relevant to pharmaceutical, agricultural, and biotechnological industries (9–12), or are impactful in biogeochemical cycles and global climate (13–16) **(Datasets S01 and S02)**. BacNa_V_ channels are thus almost certainly subject to modulatory mechanisms that vary greatly between species. Learning how these mechanisms operate can reveal new ways to influence bacteria to our benefit. To learn more about BacNa_V_ channel modulation, we focused on the C-terminal domains (CTDs) because their primary sequences and quaternary structures are different from eukaryotic Na_V_ channels and therefore unique to prokaryotes.

**Figure 1.**
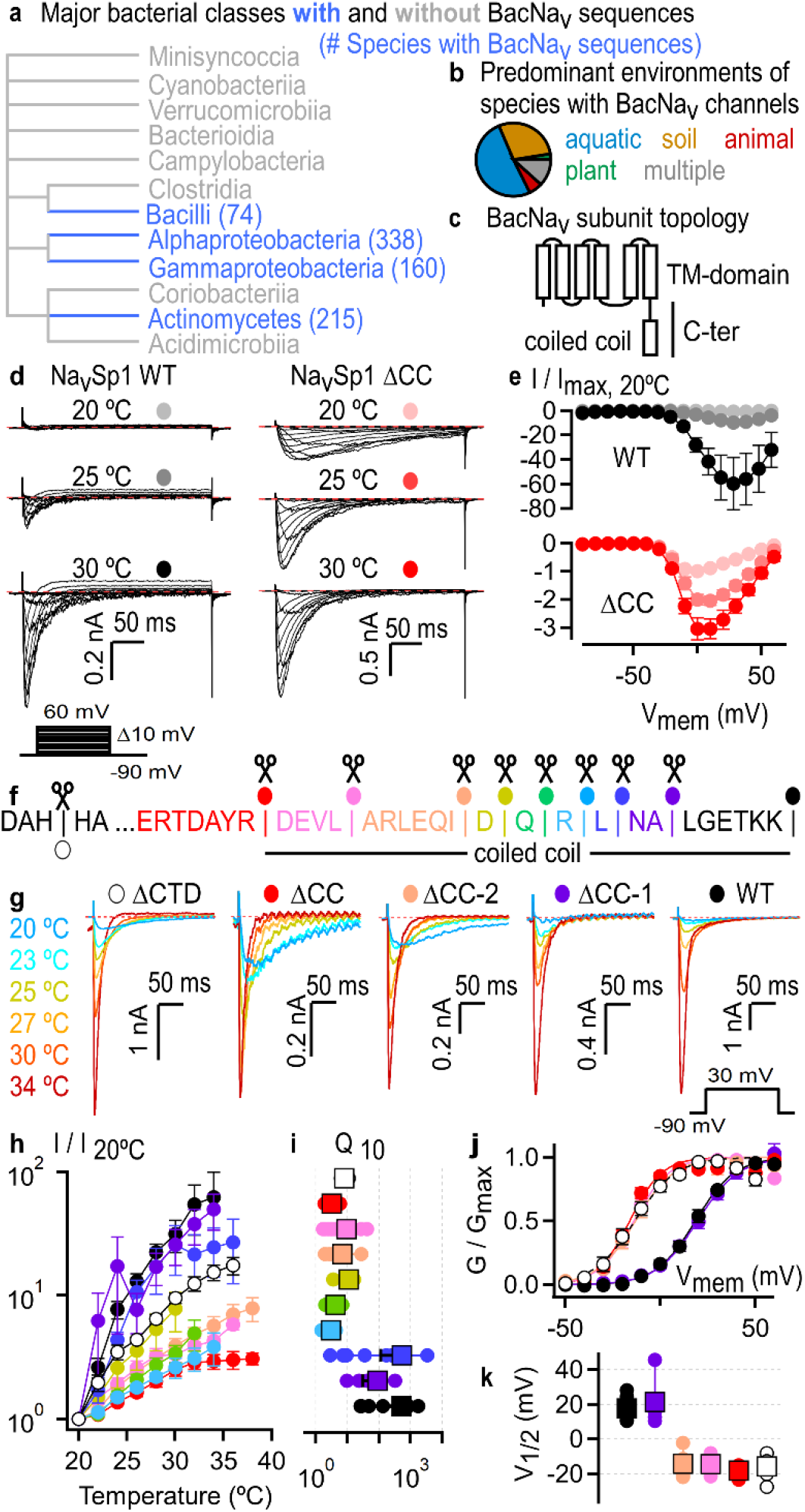
Temperature-dependent activity of Na_V_Sp1 channels requires the C-terminal coiled coil. **(a)** Taxonomy of bacterial species with BacNa_V_ channel sequences. **(b)** Predominant environments for 119 BacNa_V_-containing bacterial species. **(c)** General BacNa_V_ channel subunit topology. **(d)** Representative current families at three temperatures recorded from two cells expressing Na_V_Sp1 WT or truncated Na_V_Sp1 lacking the coiled coil. The bottom left insert illustrates the voltage-step protocol used to elicit currents. **(e)** Current-voltage relations obtained from experiments as in (d) (mean ± SEM; WT, n = 3 – 6; ΔCC, n = 4). **(f)** Amino acid sequence of the Na_V_Sp1 CTD region denoting each of the truncations that were analyzed. **(g)** Representative current traces recorded during a temperature ramp and elicited by repetitive voltage steps from −90 to 30 mV in cells expressing WT or truncated Na_V_Sp1. Current traces for only six temperatures are shown for visual clarity. **(h)** Current-temperature relations from experiments as in (g) (mean ± SEM; WT, n = 10; ΔCC-1, n = 5; ΔCC-2+DQRL, n = 8; ΔCC-2+DQR, n = 5; ΔCC-2+DQ, n = 5; ΔCC-2+D, n = 5; ΔCC-2, n = 10; ΔCC-3, n = 10; ΔCC, n = 8; ΔCTD, n = 6). **(i)** Q_10_ values from individual experiments (circles) as in (g) and their mean ± SEM (squares). **(j)** Conductance-voltage relations at room temperature obtained at 20-21 °C from recordings as in (d) (mean ± SEM; WT, n = 9; ΔCC-1, n = 8; ΔCC-2, n = 6; ΔCC-3, n = 5; ΔCC, n = 6; ΔCTD, n = 6). Continuous curves in (c) and (e) are fits to the Boltzmann equation. **(k)** V_1/2_ values from individual experiments (circles) at 20-21 °C as in (d) and their mean ± SEM (squares).

Although extremely few BacNa_V_ channels have been studied experimentally **(Dataset S03)**, the primary sequences of several of these channels are consistent with the presence of a coiled coil domain in the distal CTD (2) **(Fig. 1c)**. The CTD was not resolved in most experimentally determined structures of BacNav channels, but those where it was resolved confirm the presence of a distal coiled coil **(*SI Appendix,* Fig. S1)**. However, the conformation of the membrane-proximal region of the CTD varied greatly between experimental structures from distinct bacterial species that also represent different states of the pore –open or closed **(*SI Appendix,* Fig. S1)**. For instance, in a structure of Na_V_Ms from *Magnetococcus marinus* in the open state the proximal CTD adopts a random coil conformation (17–22) **(*SI Appendix,* Fig. S1)**. This is consistent with DEER measurements and molecular dynamics (MD) simulations of Na_V_Ms (21) that indicate that the proximal CTD is flexible. In contrast, the proximal CTD of Na_V_Ae1 from *Alkalilimnicola ehrlichii* forms a straight four-helix bundle in structures of the closed state of the channel (17, 18) **(*SI Appendix,* Fig. S1)**. The rigid character of the CTD of Na_V_Ae1 is thought to be responsible for preventing channel opening when expressed in a heterologous system (17, 18). It is unclear whether Na_V_Ae1 channel opening involves a helix to coil transition in the proximal CTD, as observed in the open-state structure of Na_V_Ms. Such a large change in secondary structure would imply a large energetic barrier for Na_V_Ae1 channel activation, potentially requiring additional factors present in the native bacterial environment that can destabilize the proximal CTD so that the pore can open. In the case of Na_V_Sp1 from the marine bacterium *Ruegeria pomeroyi* (previously known as *Silicibacter pomeroyi* (23)), a proposed temperature-driven helix to coil transition of the proximal CTD has been proposed to be required for pore opening, while also conferring heat-sensitivity to the activity of this protein (17). Interestingly, *Ruegeria pomeroyi* is an abundant and ubiquitous marine microorganism that has an important role in cloud formation by metabolizing dimethylsulfoniopropionate, an organic compound produced by marine algae, into dimethyl sulfide (24), a gas that can nucleate water droplets in the atmosphere (25, 26). The modulation of Na_V_Sp1 by temperature could be consequential for activity of *Ruegeria pomeroyi* in a warming coastal environment.

Here we used electrophysiology and computational approaches to understand the molecular mechanism through which the CTD of Na_V_Sp1 influences temperature- and voltage-dependent channel activity arising from within the transmembrane domain. We found that amino acid sequence alterations introduced into the coiled coil unleash the proximal CTD to form open state-stabilizing interactions with sites in the transmembrane domain, which in turn abolish temperature-dependent responses and favor voltage-dependent channel activation. We demonstrate that key structural determinants associated with this mechanism can be predicted with surprising accuracy by AlphaFold2 (27), which we exploit to gain insight into the structural features of the CTD in more than 800 other species of bacteria. We report that despite the low sequence identity of the CTDs from different BacNa_V_ channels, there is likely high structural conservation in this region between most channels, such that the mechanism that we described for Na_V_Sp1 is relevant in a wide number of bacterial species.

## Results

### The C-terminal coiled coil is important for Na_V_Sp1 temperature sensitivity

To investigate the role of the CTD in BacNa_V_ channel function, we focused on Na_V_Sp1 because its membrane-proximal C-terminus is proposed to function as a temperature-sensor (17). Expression of wild-type (WT) Na_V_Sp1 channels in HEK293 cells yielded robust, voltage-activated Na^+^ currents with kinetics and voltage-dependence consistent with previous observations (17, 18) **(Fig. 1d, e**, light gray). We found that heating caused a steep increase in the Na^+^ current amplitude **(Fig. 1d-h**, black) but had a relatively minor effect on the voltage-dependence of channel activation **(*SI Appendix,* Fig. S2a, b)**. Surprisingly, we found that Na^+^ currents from channels lacking the entire C-terminus (Na_V_Sp1 ΔCTD) **(Fig. 1f**, open circle) retained significant heat-sensitivity **(Fig. 1g-i**, open circles) and activated at more negative voltages (i.e. displayed a ‘left shift’ in activation voltage) than WT **(Fig. 1j, k**, open circles). In contrast, we found that channels that contained the proximal portion of the CTD but lacked the distal coiled coil (Na_V_Sp1 ΔCC) **(Fig. 1f**, red circle) lacked heat-sensitivity **(Fig. 1d-h**, red) but had similarly left-shifted voltage-dependence of activation as channels lacking the entire CTD (**Fig. 1j, k**, red). We observed a temperature-sensitivity similar to WT in channels where up to eight C-terminal amino acids were deleted **(Fig. 1f-i**, purple and blue), but truncations of nine or more residues caused a sharp reduction in heat-sensitivity **(Fig. 1f-i**, cyan, green, lime, orange, pink; ***SI Appendix,* Fig. S2e)** and a hyperpolarizing shift in the voltage-dependence of activation **(Fig. 1j, k**, orange and pink). These observations indicate that a major portion of the temperature sensitivity of Na_V_Sp1 originates in the transmembrane domain, and that the presence of the proximal CTD suppresses temperature-dependent responses unless it is connected to a distal coiled coil of least 13 amino acids in length.

Because of the lack of structures of any temperature sensitive BacNa_V_, we turned to AlphaFold (27) to build models of Na_V_Sp1 to provide structural insights into this channel and to help us to generate hypotheses about how the CTD might contribute to the temperature-sensitivity of the channel. The top-ranked model for WT Na_V_Sp1 had overall high per-residue confidence **(Fig. 2a)** and was stable in MD simulations that were run using a temperature of 50 °C **(Fig. 2b, c)**. This model was in an open state, as judged both by analysis of the structure itself **(*SI Appendix,* Fig. S3a)** and by the MD simulations in which sodium ions were seen to pass through the pore **(Fig. 2b and *SI Appendix,* Fig. S3c)**. In addition, we found that a chloride ion readily bound to a site within the coiled coil of the top-ranked model of Na_V_Sp1 that remained occupied by chloride during most of the simulation **(*SI Appendix,* Fig. S3d)**. This site formed by the side chains of R236 is analogous to a chloride binding site found in a structure of the isolated pore domain of Na_V_Ae1 (17). In relation to the state of the rest of the models, we also found both conducting (i.e. open) and non-conducting (i.e. closed or inactivated) states amongst the 24 additional models generated by AlphaFold for WT Na_V_Sp1 **(*SI Appendix,* Fig. S3b)**. Importantly, the proximal CTD had a consistent four-helix bundle structure in all 25 models **(*SI Appendix,* Fig. S3e)**, suggesting that in contrast to Na_V_Ms (17–22), drastic changes in secondary structure within the proximal CTD are not required for Na_V_Sp1 pore opening.

**Figure 2.**
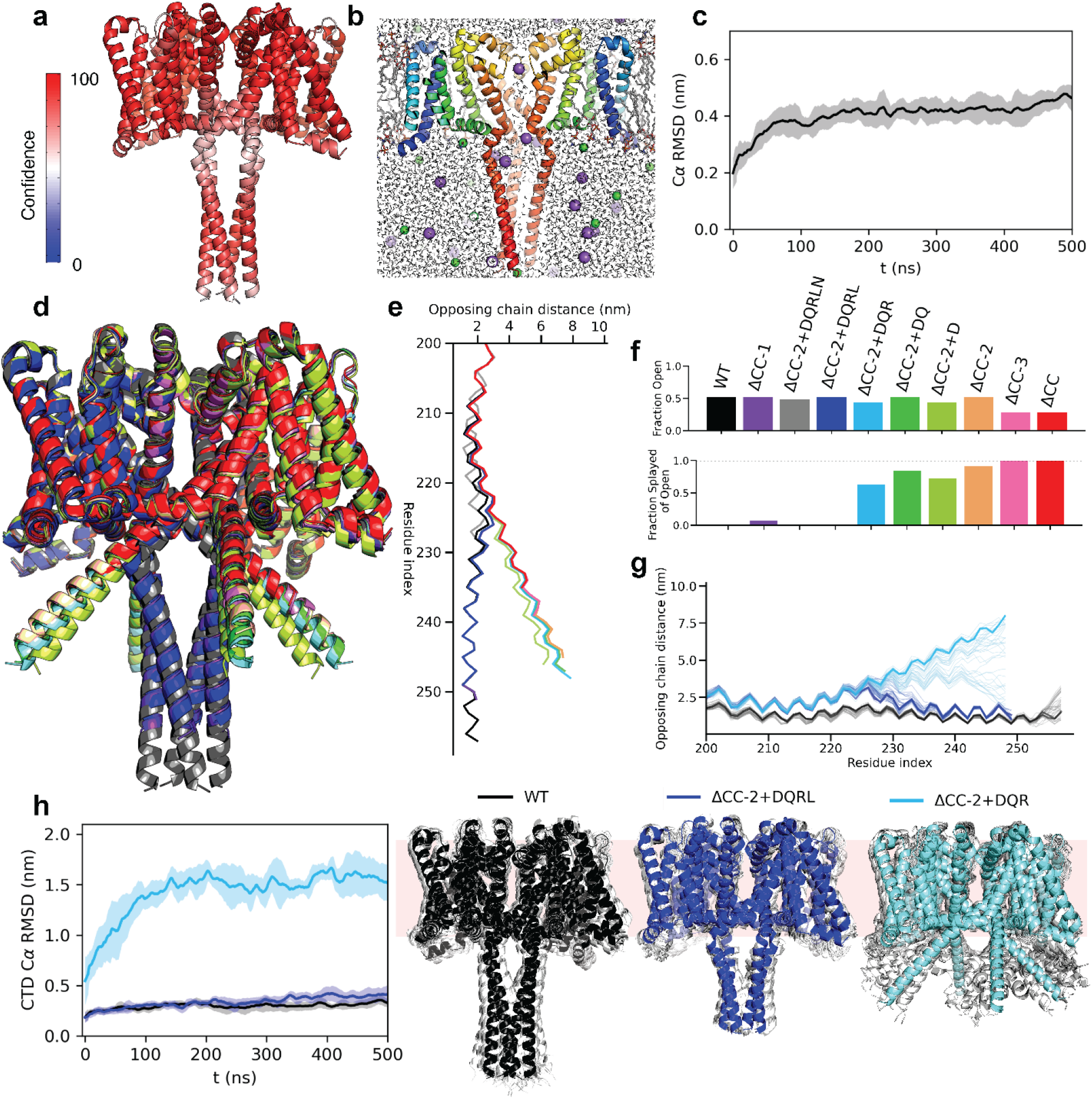
AF2 models indicate temperature-dependent activity of Na_V_Sp1 channel requires the C-terminal coiled coil. (a) Model confidence plotted on the top-scored Na_V_Sp1 AF2 model. **(b)** Representative snapshot of all-atom MD simulation of this model showing explicit membrane and sodium ions (purple) going through the pore. **(c)** Alpha-carbon RMSD of this model during the simulation represented as a sliding-window average of the three simulation replicates. **(d)** Top-ranked Alphafold2 models for full-length and truncated Na_V_Sp1 constructs, color coded as in (f). **(e)** Plots of opposing chain distance for all C-terminal residues in the top-ranked AlphaFold2 models, color coded as in (f). **(f)** Top: Fraction of models with an open pore out of all 25 AF2 models generated per construct, as determined by the distance between opposing subunits at position I216. Bottom: Fraction of models in a splayed conformation out of the open models per construct, as determined by the distance between opposing subunits at position T232. **(g)** Opposing subunit per-residue distances along the CTD of WT (black), ΔCC2+DQRL (truncation of eight amino acids, dark blue), and ΔCC2+DQR (nine truncated amino acids, light blue) for the first 400 ns of simulation. The flexibility of the splayed helices in ΔCC2+DQR is clearly visible**. (h)** RMSD of the CTD of WT (black), ΔCC2+DQRL (dark blue), and ΔCC2+DQR (light blue) represented as a sliding-window average of the three simulation replicates. The right panel shows the initial models in color overlaid with structural snapshots (white) captured every 20 ns during a single replicate of the simulations for each construct.

Next, we built AlphaFold models for each of the truncated Na_V_Sp1 constructs whose temperature-sensitivity we had measured, and found a strong correlation between the structure of the CTD in the models and the experimentally-determined temperature sensitivity of the channels: 99 out of 100 models of channels that had strong temperature-sensitivity had a helical bundle CTD fold closely resembling WT Na_V_Sp1 **(Fig. 2d-f**, black, purple, gray, blue), while the C-terminal helix bundles were completely splayed in the majority of the models for each of the channels that had reduced temperature sensitivity **(Fig. 2d-f**, light blue, green, lime, orange, pink, red). Importantly, the fraction of models representing open and non-conducting states was similar across the two groups of channels **(Fig. 2f**, top), indicating that the predicted closed or open state of the pore did not determine whether the CTD was splayed or not in the models. We then conducted MD simulations using the top-ranked models of truncated Na_V_Sp1 channels with either strong (ΔCC-2+DQRL, blue) or weak (ΔCC-2+DQR, cyan) temperature-sensitivity to test the stability of the structures predicted by AlphaFold. We ran simulations at a temperature of 50 °C because it resulted in an appropriate membrane lipid fluidity while accelerating conformational sampling during the simulations and providing a more stringent test on model stability. We observed that the CTDs remained stable during simulations for both WT and the strongly temperature-dependent ΔCC-2+DQRL channels **(Fig. 2g, h**, blue and black), while dynamics were substantial within the splayed CTDs of the ΔCC-2+DQR channels that had reduced temperature dependence **(Fig. 2g, h**, cyan). Our results so far suggest that Na_V_Sp1 requires the integrity of the distal coiled coil to have strong sensitivity to heat and that truncations greater than eight amino acids can cause the coiled coil to disassemble at room temperature, resulting in loss of temperature sensitivity.

Having found that coiled coil truncations of a certain length lead to loss of temperature sensitivity, we tested the possibility that temperature-dependent splaying of the Na_V_Sp1 coiled coil could be associated with the response of the channel to heat. To this end, we examined whether formation of inter-subunit disulfide bonds under oxidizing conditions between a pair of cysteines substituted into the coiled coil (***SI Appendix,* Fig. S4a)**, which should prevent it from splaying, would influence channel activity and heat sensitivity. Using a circularly permuted HaloTag (28) fused to the N-terminus of Na_V_Sp1, we were able to fluorescently label channels expressed in cells to then detect cysteine-crosslinked channel complexes after non-reducing SDS-PAGE of the cell lysate (***SI Appendix,* Fig. S4b)**. We found that crosslinked channel tetramers predominated in cells that were exposed to oxidation with 40 mM H_2_O_2_ (***SI Appendix,* Fig. S4b, c**; blue), while monomers predominated in untreated cells (***SI Appendix,* Fig. S4b, c**; black) or in cells exposed to a lower concentration of H_2_O_2_ (***SI Appendix,* Fig. S4b, c**; cyan). Importantly, we confirmed that inter-subunit crosslinking was reversed by DTT **(*SI Appendix,* Fig. S4b, c**; beige and brown), and that crosslinking occurred exclusively between the two cysteines that we substituted into the coiled coil because we did not detect any crosslinked complexes in HaloTag channel-fusions lacking the two substituted cysteines **(*SI Appendix,* Fig. S4b**, left). Having established that exposure to H_2_O_2_ increases the fraction of crosslinked tetramers in cells, we tested whether this treatment affected channel responses to heat or voltage. Whole-cell currents mediated by cysteine-substituted channels after exposure to H_2_O_2_ had similar temperature- and voltage-sensitivity as WT channels **(*SI Appendix,* Fig. S4d, e)**. To rule out the possibility that the currents measured after H_2_O_2_ treatment were generated exclusively by non-crosslinked channels while cysteine-crosslinked channel complexes remained silent, we assessed the current magnitude in H_2_O_2_-exposed cells before and during exposure to the reducing agent DTT **(*SI Appendix,* Fig. S4f)** and observed no evidence of DTT-induced current recovery. Together, our results establish that coiled coil disassembly is not part of the activation mechanism of Na_V_Sp1, and that a stably folded coiled coil is likely required for Na_V_Sp1 channels to respond to temperature with high sensitivity

### Coupling between the distal and proximal regions of the C-terminus determines temperature sensitivity of Na_V_Sp1

Our findings so far are consistent with the view that the C-terminal coiled coil contributes to channel function as a rigid scaffold for the rest of the CTD, and so its specific sequence should bear no influence on channel activity so long as it folds as a coiled coil. To test this hypothesis, we generated chimeric Na_V_Sp1 constructs containing coiled coil sequences of BacNa_V_ channels from other species of bacteria while keeping the rest of the Na_V_Sp1 sequence unchanged **(Fig. 3a)** and measured their sensitivity to temperature **(Fig. 3b)** and voltage **(Fig. 3c)**. Three of the channels we selected for their distal C-terminus coiled coil sequences had been previously characterized at the structural and/or functional level (Na_V_Bh (5, 29, 30), Na_V_Ab1 (31), Na_V_Ms (19–21)) **(*SI Appendix,* Fig. S1, Dataset S03)**, while two others (Na_V_Rbl, Na_V_Ob) are uncharacterized, but their distal CTD primary sequences differ markedly from one another in their probabilities of folding as a coiled coil **(*SI Appendix,* Fig. S5a)**. Contrary to the hypothesis, we found that only one of the chimeras (Rbl, teal) had WT-like temperature-sensitivity, while the rest of the chimeras had either moderate (Ob, purple; Bh, blue) or weak (Ms, yellow; Ab1, gray) temperature-sensitivity similarly to channels lacking their coiled coil region (ΔCC, red) **(Fig. 3b, f and *SI Appendix,* Fig. S2e)**. Interestingly, we found that the voltage-sensitivity of the chimeras tended to be correlated with their temperature-sensitivity **(Fig. 3c, g)**, whereby the two chimeras with the weakest temperature-dependence, Ms and Ab1, had the largest hyperpolarizing shifts in their voltage of activation, similarly to the voltage-dependence of ΔCC channels.

**Figure 3.**
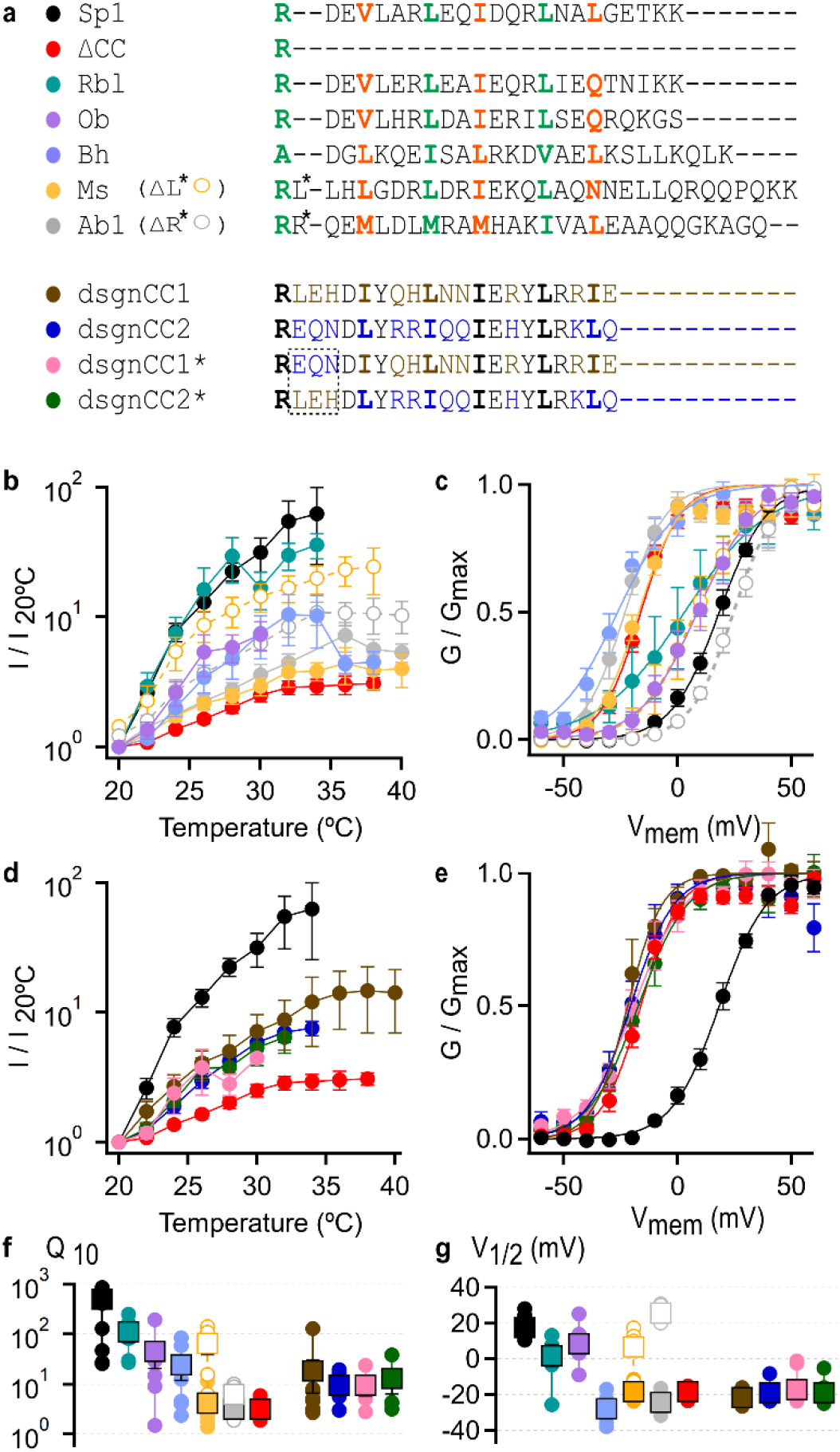
The coiled coil sequence strongly influences the temperature-dependence of Na_V_Sp1 channels. **(a)** Na_V_Sp1 coiled coil-chimera amino acid sequence alignment. Coiled coil heptad positions are highlighted in green and orange. **(b)** Current-temperature relations for chimeras with coiled coils from other BacNa_V_ channels (mean ± SEM; Rbl, n = 5; Ob, n = 7; Bh, n = 7; Ms, n = 10; Ms ΔL248, n = 5; Ab1, n = 9; Ab1 ΔR248, n = 5). **(c)** Conductance-voltage relations at room temperature for chimeras with coiled coils from other BacNa_V_ channels (mean ± SEM; Rbl, n = 6; Ob, n = 6; Bh, n = 7; Ms, n = 9; Ms ΔL248, n = 5; Ab1, n = 9; Ab1 ΔR248, n = 5). **(d)** Current-temperature relations for chimeras with synthetic coiled coils (dsgnCC1, n = 11; dsgnCC2, n = 8; dsgnCC1*, n = 5; dsgnCC2*, n = 5). **(e)** Conductance-voltage relations at room temperature for chimeras with synthetic coiled coils (dsgnCC1, n = 3; dsgnCC2, n = 4; dsgnCC1*, n = 9; dsgnCC2*, n = 6). **(f)** Q_10_ values from individual experiments (circles) as in (b) and (d) and their mean ± SEM (squares). **(g)** V_1/2_ values from individual experiments (circles) as in (c) and (e) and their mean ± SEM (squares). Continuous curves in (c) and (e) are fits to the Boltzmann equation.

Alignment of the coiled coil primary sequences based on their heptad-repeat motifs revealed that the sequences of the two chimeras with the weakest temperature dependence, Ab1 and Ms, had a single-residue insertion at the junction between the membrane-proximal portion of the CTD and the N-terminal end of the coiled coil region **(Fig. 3a)**. We found that removal of this extra residue (Ms ΔL, yellow open circles) increased the temperature-dependence of the Ms chimera almost to the same level as WT channels **(Fig. 3b, f)**, which was also accompanied by a depolarizing shift in voltage-dependence **(Fig. 3c, g)**. The single residue deletion also caused an increase in temperature-sensitivity of the Ab1 chimera (Ab1 ΔR, gray open circles) that was smaller than for Ms **(Fig. 3b, f)**, and caused a greater depolarizing shift in voltage sensitivity **(Fig. 3c, g)**. These findings reveal that the temperature- and voltage-dependent activity of Na_V_Sp1 channels is highly sensitive to the amino acid sequence of the distal CTD.

The distal CTD sequences that resulted in the lowest temperature-sensitivity have been experimentally found to fold as a coiled coil in Na_V_Ab1 (31) and Na_V_Ms channels (19, 21), so it is unlikely that their decreased sensitivity to temperature was caused by coiled coil misfolding. To further address this possibility, we used Rosetta (32, 33) to design *de novo* two sequences predicted to fold as a tetrameric coiled coil (**Fig. 3a**, dsgnCC1 and dsgnCC2) and measured the temperature- and voltage-sensitivity of Na_V_Sp1 chimeras containing these sequences in place of their native distal CTD sequences. The voltage dependence of the two chimeras containing *de novo*-designed sequences was shifted towards negative voltages as observed in ΔCC channels lacking a coiled coil (**Fig. 3e, g**). The temperature sensitivity in both chimeras was considerably weaker than WT but higher than in ΔCC channels (**Fig. 3d, f**; brown, marine), most closely resembling Bh in both temperature-sensitivity and voltage-dependence. We observed the same temperature-sensitivity and voltage-dependence in two additional chimeras where the first three amino acids in the distal C-terminus sequence were swapped **(Fig. 3a, d-g**, dsgnCC1*, pink; dsgnCC2*, green), conforming that all designed chimeras had a similar effect on channel function regardless of their specific sequences. Together, our findings so far establish that even small differences in the primary sequence of the distal CTD can have profound effects on the temperature- and voltage-sensitivity of Na_V_Sp1 channels even though all the distal CTD sequences that we examined are likely to fold as tetrameric coiled coils.

To generate hypotheses about the structural mechanism by which changes in the sequence of the distal CTD can influence Na_V_Sp1 channel activity, we used AlphaFold2(27) to generate 25 structural models for each of the coiled coil chimeras. The distal CTD adopted a similar coiled coil structure in all models, yet the structure of the proximal CTD showed marked differences relative to WT in the top-scored models for the Ms and Ab1 chimeras that had the weakest temperature sensitivity **(Fig. 4a, b**; gray and yellow). Specifically, these models had a helical kink centered around residue E227 in the membrane-proximal CTD, eight amino acids away from the N-terminal end of the coiled coil region **(Fig. 4b)**, which contrasted with the straighter helix bundle configuration in WT or the other chimeras **(Fig. 4b**; black, blue, purple, and teal). To assess these differences between each of the 25 models that we generated per chimera, we mapped each model on a 2D plot describing the pore diameter on the x-axis to distinguish between closed and open state models, and the distance between opposing subunits at E227 on the y-axis to distinguish models with a kinked helix bundle in their proximal CTD **(Fig. 4c)**. We obtained models with open or closed pores for each of the chimeras. Interestingly, for the Ab1 and Ms chimeras that had the weakest temperature-sensitivity, both open- and closed-pore models were clustered away from the rest of the models based on the inter-subunit distance at position E227 **(Fig. 4c**, gray and yellow). All models of the Ab1 and Ms chimeras with an open pore presented a kink at E227 **(Fig. 4e, bottom)**, while the proximal CTD had a straighter four-helix bundle structure in most or all open state models for WT and the rest of the chimeras with stronger temperature sensitivity, including the single-residue deletions Ms ΔL* and Ab1 ΔR* (**Fig. 4d, e**). Interestingly, all open-state models of chimeras containing *de novo* designed sequences had a kink at position E227 (**Fig. 4e**), consistent with all other variants showing reduced temperature-sensitivity compared to WT.

**Figure 4.**
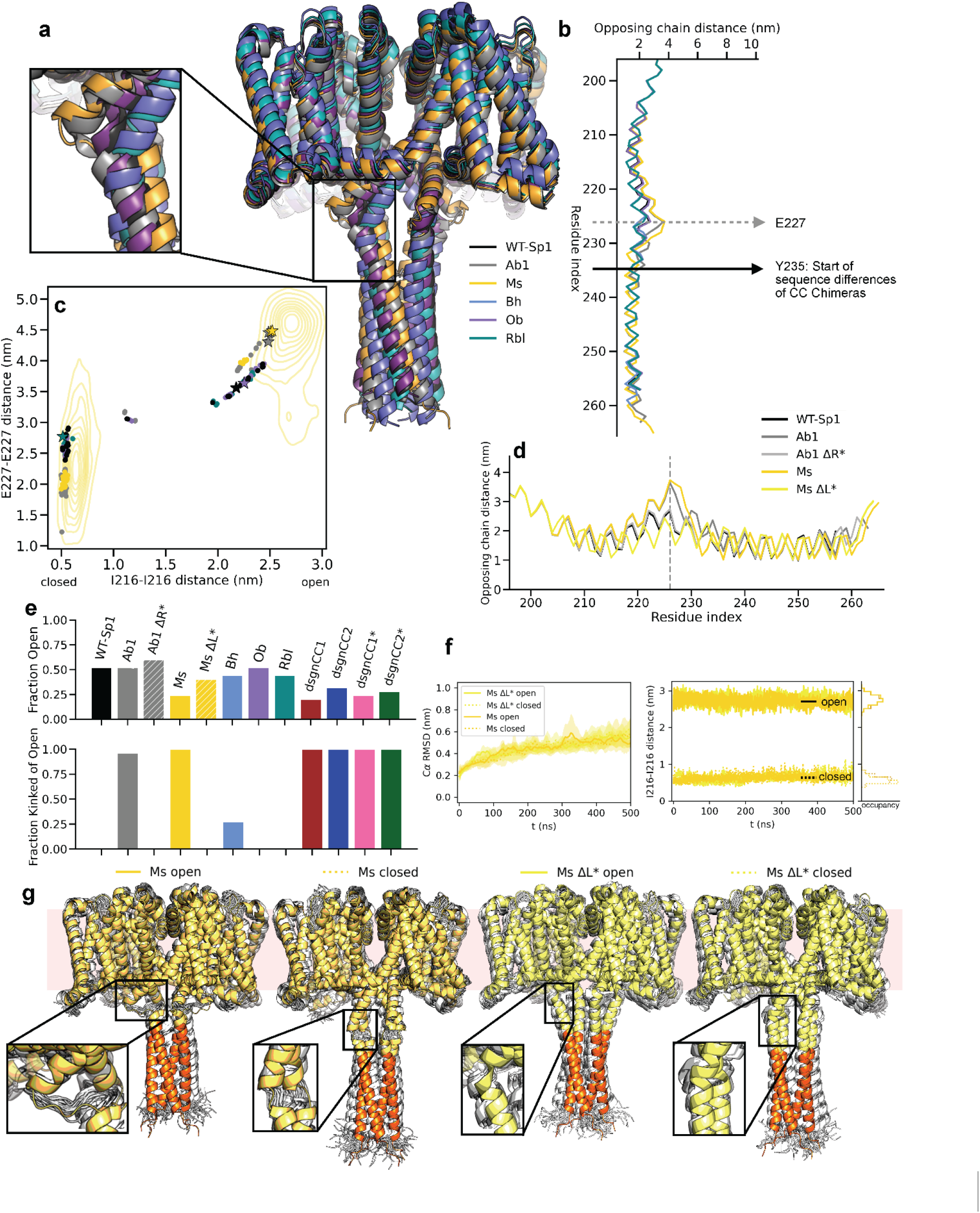
The coiled coil sequence strongly influences the conformation of Na_V_Sp1 channels. **(a)** Top-ranked AlphaFold models of all chimeras overlayed: black – WT Na_V_Sp1, grey – Ab1, gold – Ms, light blue – Bh, purple – Ob, and teal – Rbl. **(b)** Opposing chain distances along the CTD for the top-ranked models for all chimeras. A kink at position E227 is seen in the Ab1 (grey) and Ms (gold) chimera models. **(c)** Plotting all 25 AF2 models based on their E227-E227 distance between opposing subunits, which is sensitive to the presence of a kink, and the I216-I216 distance between opposing subunits, which is sensitive to the open or closed state of the pore. Data for the Bh chimera models were omitted for clarity. Stars indicate top ranked models. Contours indicate the density of states sampled during MD simulations of the Ms chimera in closed or open pore states. **(d)** Opposing chain distances along the CTD for the top-ranked models of Ms (gold), Ms ΔL (yellow), Ab1 (grey), and Ab1 ΔR (light grey). **(e)** Top: Fraction of models with an open pore out of all 25 models obtained per chimera as determined by the distance between opposing subunits at position I216. Bottom: Fraction of open models with a kink at E227 out of all 25 models obtained per chimera. **(f)** Alpha-carbon RMSD (left) and opposing chain distance of I216 (right) during simulations for the top-ranked models for Ms and Ms ΔL started from a closed or open pore state. RMSD (left), shown as a sliding-window average across all three replicates, confirms that all four types of structures are equally stable. The I216 distances (right), shown as the time-dependent trace for a single simulation for each construct, confirm that during the entire length of the simulations the pores of each of these constructs remained either closed or open, depending on the state from which simulations were started. The right-most inserts show histograms for the I216 distance from all three 500 ns replicates. For the open states, the Ms ΔL histogram has a mean = 2.7568 nm and a Standard deviation (std) = 0.13, and Ms has a mean = 2.7429 nm and std = 0.13 nm. For the closed states, Ms ΔL has mean = 0.5702 nm with std = 0.08 nm and Ms has mean = 0.5873 nm with std = 0.09 nm. **(g)** Top-ranked models with closed or open pores of Ms and Ms ΔL from which simulations were started, shown in gold or yellow, respectively, with the coiled coil colored in orange. Structural snapshots at every 20 ns of the simulation are shown in white.

Our observations so far suggest that perturbations in the sequence of the coiled coil in the distal CTD of Na_V_Sp1 propagate into the structure of the proximal CTD, which may allow this region to adopt a kinked conformation in the open state of the channel. To determine whether the AlphaFold models having a kinked proximal CTD represented stable structures, we carried out MD simulations focusing on the Ms and Ms ΔL* chimeras because their sequences differ by just one amino acid while showing striking differences in their temperature-sensitivity. We started simulations from the top ranked models with closed and open pores for each construct, all of which produced stable MD simulations at 50 °C showing no pore gating transitions or major changes in the structure of the CTD **(Fig. 4f, g)**. Importantly, we observed no kink formation within the proximal CTD during the simulations of the Ms ΔL* chimera in the open state, indicating that the predicted straighter four-helix bundle arrangement of the Ms ΔL* chimera and the kinked proximal CTD of Ms both represent stable protein conformations.

Together, our findings indicate that perturbations to the coiled coil in the distal CTD can affect the conformation of the rest of the channel in a mechanism that is likely to involve formation of a kink within the proximal CTD in the open state that is somehow associated with a reduction in temperature-dependent channel activity. To gain insight into why the amino acid sequence of the coiled coil may influence the structure of the helical bundle in the proximal CTD, we analyzed the geometry of the coiled coil region in each of the top-ranked chimera models using a program that measures the deviation of a given helical bundle structure from an idealized coiled coil (34). We found that the axial rotation of the individual helices relative to the central axis of the coiled coil **(*SI Appendix,* Fig. S5b)** differed more markedly at the N-terminus of the coiled coil between channels with the lowest temperature-sensitivity (Ab1, Ms, dsgnCC1, dsgnCC2, dsgnCC1*, dsgnCC2*) relative to the rest (WT, Rbl, Ob, Bh, Ms ΔL, Ab1 ΔR) **(*SI Appendix,* Fig. S5c, d)**. This suggests that differences in the axial rotation at the N-terminal end of the coiled coil helices between the distinct chimeras could influence the structure of the proximal CTD by modifying the tension that this region experiences from laying between two rigid helical bundles – the S6 helices and the coiled coil.

### Tuning of Na_V_Sp1 activity through interactions between the C-terminus and the transmembrane domain

Our results so far suggest that loss of temperature-dependent activity by the Na_V_Sp1 channel is associated with kinking of the four-helix bundle in the proximal CTD, which in turn can be caused by removal or perturbation of the coiled coil at the distal CTD. To determine the mechanism through which the membrane-proximal CTD can suppress temperature-dependent Na_V_Sp1 activity, we inspected the AlphaFold models for evidence of interactions formed by the proximal CTD. Indeed, in the top ranked open state models for the Ab1 and Ms chimeras with kinked proximal CTDs **(Fig. 5a**, gray and yellow), as well as for truncated Na_V_Sp1 models with splayed CTDs **(Fig. 5c, d**, cyan and red), we found that residues E227 in the proximal CTD and R117 in the S4-S5 linker helix in the transmembrane domain were sufficiently close to interact, while the same two residues were too far to form interactions in the top ranked open state model of WT Na_V_Sp1 **(Fig. 5a**, black). Interactions between equivalent conserved positions in the proximal C-terminus and the S4-S5 linker have been previously described in structures of the open state of Na_V_Ms (19), and these interactions were proposed to play an important role stabilizing the open state of the Na_V_Ms channel (35, 36). Importantly, these same interactions were also shown to stabilize the open state of Na_V_Sp1 at room temperature (37).

**Figure 5.**
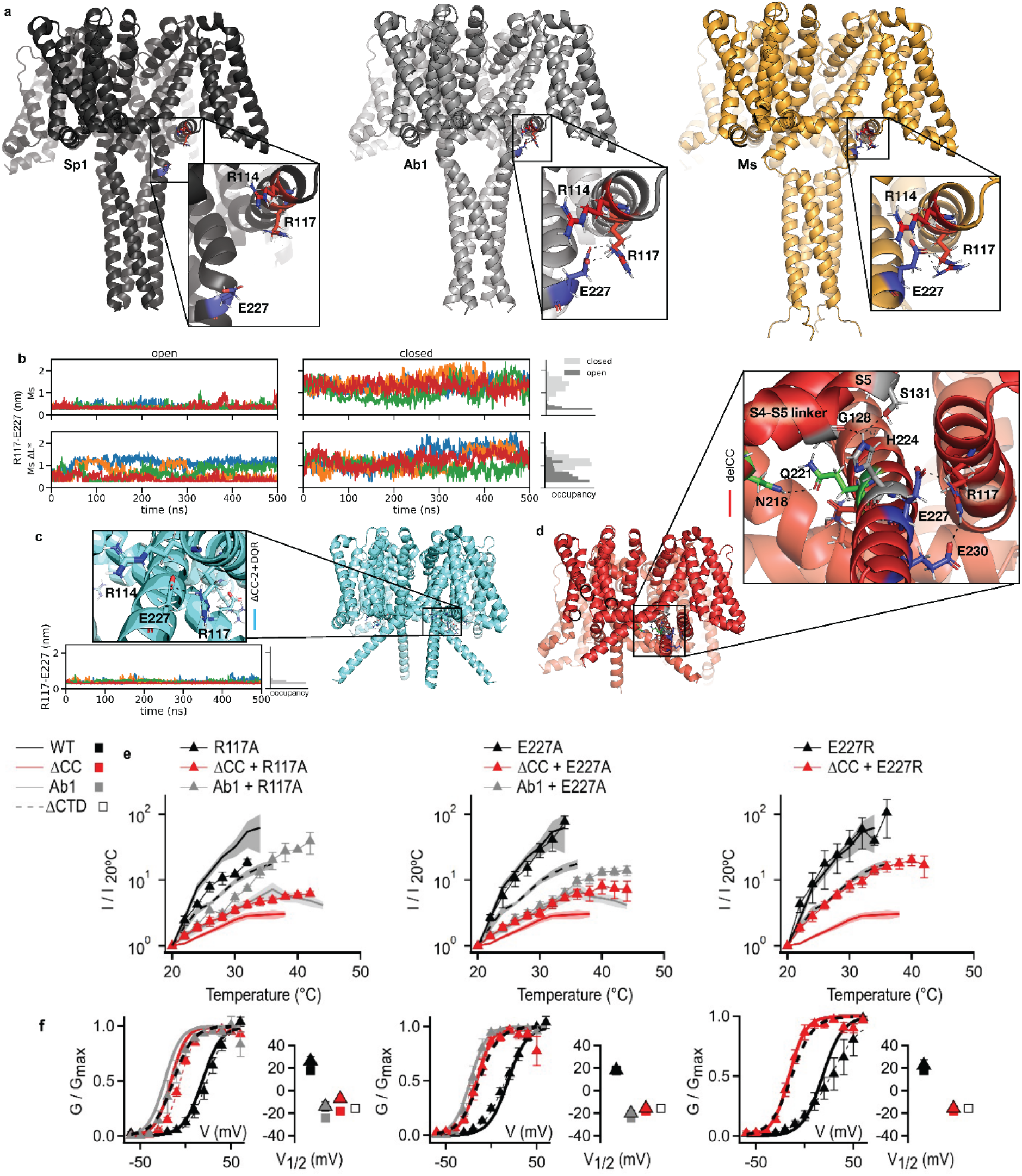
Interactions between the C-terminus and the S4-S5 linker tune Na_V_Sp1 activity. **(a)** Top-ranked models of select channel constructs (black – WT Na_V_Sp1; grey – Ab1 chimera; gold – Ms chimera). Interactions between E227, R117, and R114 (shown as sticks) are shown. **(b)** Closest heavy atom distance between E227 and R117 shown for a single simulation started from the open (left) or closed (right) pore states of the Ms and Ms ΔL chimeras. The insert on the right shows the histograms for the distances measured over all three simulation replicates for each channel in the open or closed state. For Ms the open state mean = 0.3345 and std = 0.09 and the closed state mean = 1.3342 and std = 0.29. For Ms ΔL the open state mean = 0.6400 and std = 0.32 and the closed state mean = 1.1089 and std = 0.28. In both cases the open state has E227 and R117 more consistent with interaction range. **(c, top)** Top-ranked model for ΔCC-2+DQR (cyan) with a splayed CTD. **(c, bottom)** Closest heavy atom distance between E227 and R117 during a single simulation started from the open state of ΔCC-2 +DQR, with a histogram on the right showing the distribution of this distance across all three replicates (mean = 0.3529, std = 0.1105). **(d)** Top-ranked model for ΔCC (red) with a splayed C-terminus, showing additional interactions between the proximal CTD and the S4-S5 linker. **(e, left)** Normalized current-temperature relations measured at 30 mV (mean ± SEM; WT, black continuous curve, n = 10; ΔCC, red continuous curve, n = 8; Ab1, gray continuous curve, n = 9; ΔCTD, dashed curve, n = 6; Na_V_Sp1 R117A, black triangles, n = 4; ΔCC R117A, red triangles, n = 5; Ab1_cc_ R117A, gray triangles, n = 5). **(e, middle)** Normalized current-temperature relations measured at 30 mV (mean ± SEM; Na_V_Sp1 E227A, black triangles, n = 5; ΔCC E227A, red triangles, n = 5; Ab1 E227A, gray triangles, n = 6). **(e, right)** Normalized current-temperature relations measured at 30 mV (mean ± SEM; Na_V_Sp1 E227R, black triangles, n = 5; ΔCC E227R, red triangles, n = 7). **(f, left)** Conductance-voltage relations obtained at room temperature (mean ± SEM; WT, black continuous curve, n= 9; ΔCC, red continuous curve, n = 6; Ab1, gray continuous curve, n = 9; ΔCTD, dashed curve, n = 6; Na_V_Sp1 R117A, black triangles, n = 3; ΔCC R117A, red triangles, n = 5; Ab1 R117A, gray triangles, n = 5). **(f, middle)** Conductance-voltage relations obtained at room temperature (mean ± SEM; Na_V_Sp1 E227A, black triangles, n = 5; ΔCC E227A, red triangles, n = 5; Ab1 E227A, gray triangles, n = 5). **(f, right)** Conductance-voltage relations at room temperature (mean ± SEM; Na_V_Sp1 E227R, black triangles, n = 5; ΔCC E227R, red triangles, n = 5). The inserts show the V_1/2_ obtained from fits to the Boltzmann equation.

We next examined our MD simulations to determine whether stable interactions between residues E227 and R117 were observed as suggested by the models. In MD simulations that were started from top-ranked open state models for the Ms or Ms ΔL chimeras **(Fig. 5b**, left), occupancy of the E227-R117 interaction was much higher for the Ms model with a kinked proximal CTD than for the Ms ΔL model that did not have a kink in the proximal CTD. Importantly, the interaction between E227 and R117 did not stably form in the simulations from models that had a closed pore **(Fig. 5b**, right). Residues E227 and R117 also showed stable, long-lived interactions in simulations of truncated ΔCC-2+DQR Na_V_Sp1 channels with a splayed CTD **(Fig. 5c)**. In the ΔCC-2+DQR splayed model simulation we observed that additional residues E230 in the proximal CTD and R114 in the S4-S5 linker helix also formed interactions, albeit with a lower occupancy than E227 and R117 **(*SI Appendix,* Fig. S6a)**. Additional interactions with similar occupancy could also be detected in the same interface involving positions H224 in the C-terminus and G128 and S131 in the S4-S5 linker of the neighboring subunit **(*SI Appendix,* Fig. S6b)**. Similar interactions are likely formed in the open state of Na_V_Sp1 ΔCC, as all residues involved are within interaction distance in the top-ranked model for this truncated channel **(Fig. 5d)**. We also tested whether running our simulations at a higher temperature changed the propensity for E227 and R117 to interact and found that although other parts of the protein were less stable, this interaction persisted even at higher temperatures **(*SI Appendix,* Fig. S6c)**.

To determine whether the interactions formed by splayed or kinked conformations of the proximal CTD and the S4-S5 linker helix are responsible for suppressing temperature-dependent Na_V_Sp1 channel activity, we investigated whether there was an effect on temperature sensitivity from mutations predicted to destabilize the E227-R117 interaction. We found that alanine substitutions at positions R117 in the S4-S5 linker **(Fig. 5e**, left; ***SI Appendix,* Fig. S7)** or E227 in the proximal CTD **(Fig. 5e**, middle; ***SI Appendix,* Fig. S7)** caused a modest but detectable increase in the response to temperature when introduced into Na_V_Sp1 ΔCC **(Fig. 5e**, red triangles) or the Ab1 chimera **(Fig. 5e**, gray triangles; ***SI Appendix,* Fig. S7)**, which in the open state are predicted to have splayed or kinked CTDs. The alanine substitutions did not increase the temperature responses when introduced into WT Na_V_Sp1 **(Fig. 5f**, black triangles; ***SI Appendix,* Fig. S7)**, and all mutations had minimal effect on voltage-sensitivity **(Fig. 5f**, left and middle; ***SI Appendix,* Fig. S7)**. We reasoned that the individual alanine substitutions at positions R117 or E227 had failed to fully restore temperature-dependent activity to the same level as Na_V_Sp1 ΔCTD channels lacking the entire C-terminus **(Fig. 5e, f**, black continuous curves) because the interface between the CTD and the S4-S5 linker is stabilized by multiple interactions involving other residues besides R117 and E227. We therefore tested the effect of reversing the charge at position E227, which we predicted would result in strong electrostatic repulsion between the CTD and the S4-S5 linker that contains multiple positively charged residues **(Fig. 5a, d)**. Consistent with our hypothesis, we found that the E227R substitution increased the temperature dependence of Na_V_Sp1 ΔCC to the same level as that of Na_V_Sp1 ΔCTD channels **(Fig. 5e**, red triangles; ***SI Appendix,* Fig. S7)** and had no effect when introduced into full-length Na_V_Sp1 **(Fig. 5e**, black triangles; ***SI Appendix,* Fig. S7)**. Based on our observations, we conclude that in the open state of Na_V_Sp1, the proximal CTD can interact with the S4-S5 linker and suppress the temperature-dependent activity of the channel when it adopts a kinked or splayed conformation.

### Structural conservation across BacNa_V_ channels suggests a common regulatory role for the C-terminus

We next wondered whether a similar mechanism as we have described for the Na_V_Sp1 CTD could regulate the function of other BacNa_V_ channels. A multiple sequence alignment from 825 non-redundant BacNa_V_ channels **(Dataset S01)** showed strong sequence conservation within the six transmembrane helices and the pore **(Fig. 6a)**, consistent with a previous sequence analysis that focused on the voltage-sensor and pore domains of diverse BacNa_v_ channels (38). In contrast, the sequence identity within the N-terminus, C-terminus, and surprisingly the S5 helix was remarkably lower than it was in the rest of the transmembrane domain. The high sequence identity within the S6 helix region allowed us to define the approximate start of the CTD for all sequences. The shortest of these channels at 218 residues had no CTD, and the longest was 365 residues long with a CTD of 100 residues; the longest CTD overall was 112 residues and the shortest CTD of any channel that had a CTD was only 7 residues long **(Fig. 6b)**. Overall, sequences could be grouped into two populations based on the length of the CTD, with a major population of shorter length that included Na_V_Sp1, and a smaller population with longer lengths. Notably, almost all BacNa_v_ channels had CTDs of 28 amino acids or longer, with only six shorter CTD sequences that were 0, 7, 9, 11, 17, and 22 amino acids long (**Fig. 6b**). Most sequences had a CTD similar in length to that of Na_V_Sp1, which is 37 amino acids long. The high variability in the length and amino acid composition of the CTD between channels from distinct species suggests that some of these differences might reflect adaptations to the specific environments of the bacteria. To gain more insight into these variations, we analyzed the amino acid content of the CTD sequences. While 402 CTDs had no proline residues, as would be expected for a predominantly α-helical domain, some outliers had as many as 23% proline residues in the CTD sequence **(Fig. 6c)**. The distribution of sequences based on the number of charged residues in the CTD was relatively symmetric, where some CTD sequences had as few as 9% charged residues while others like the CTD of the BacNa_V_ from *Pseudovibrio denitrificans* had 63% charged residues **(Fig. 6d)**.

**Figure 6.**
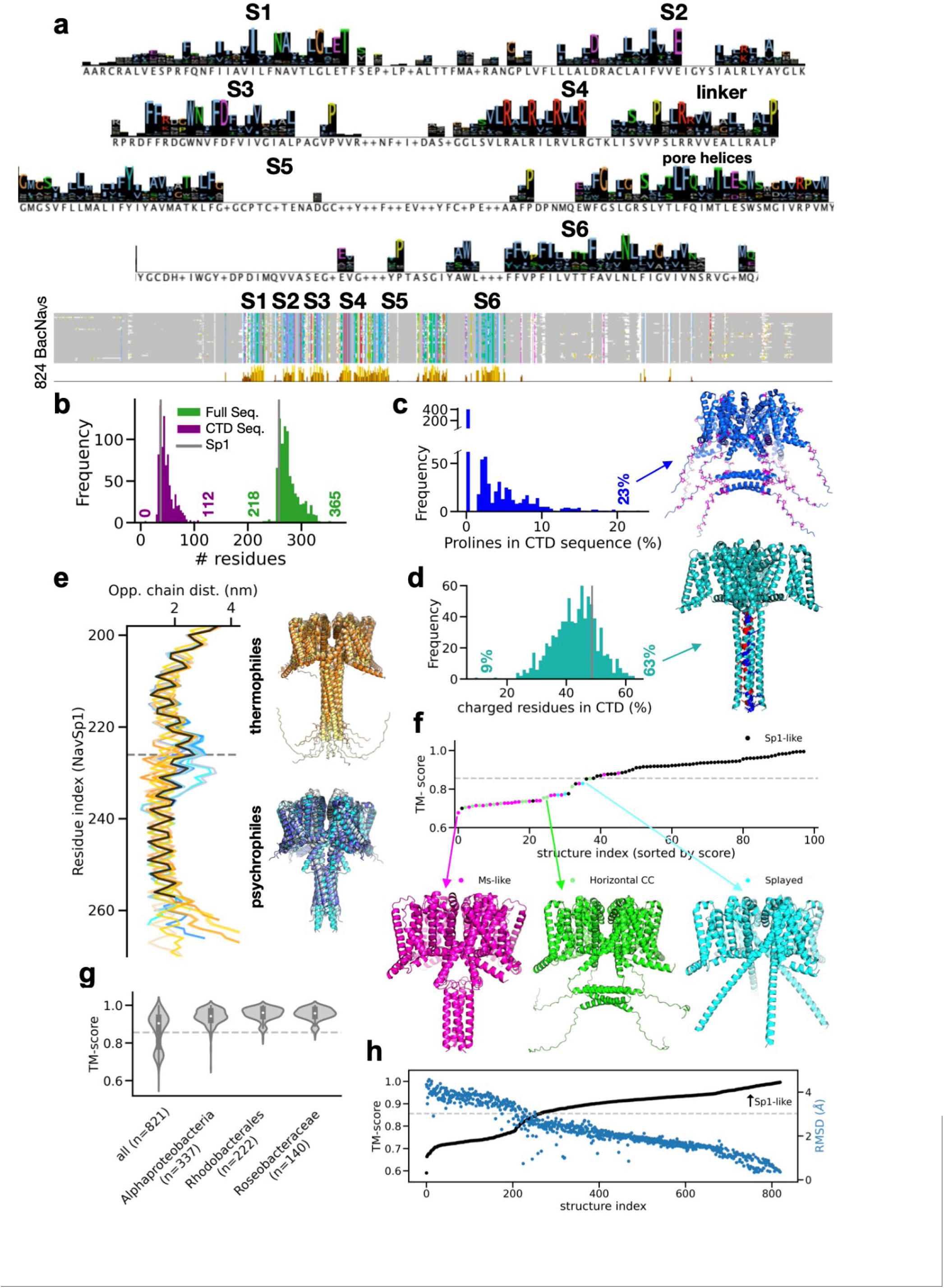
Sequence and structural conservation across BacNa_V_s channels. **(a)** WebLogo obtained from the alignment of 824 non-redundant BacNa_V_ channel sequences, as shown on the bottom panel; sequences are colored by the Clustal coloring scheme. **(b)** Histograms of BacNa_V_ channel sequences binned by the length of their CTD (burgundy) or total protein length (green). The vertical black lines denote the position of the Na_V_Sp1 sequence. **(c)** Histograms of BacNa_V_ channel sequences binned by the percentage of prolines in the CTD sequence. 402 sequences had zero prolines in the CTD, while the sequence from *Streptomyces prasinopilosus* had 23% prolines in the CTD. Here the top model is shown with prolines with the CTD and transmembrane domain shown in sticks and colored in magenta. **(d)** Histograms of BacNa_V_ channel sequences binned by the number of charged residues in the CTD. The top model for the sequence from *Pseudovibrio denitrificans*, in which 63% of its CTD residues are charged, is shown. Positive residues in the CTD or transmembrane domain are colored in blue, and negative residues in red. **(e)** Opposing chain distances along the C-terminus for the top-ranked models for sequences from five thermophilic (orange) and eight psychrophilic (blue) bacterial species in our dataset. Distances for WT Na_V_Sp1 are shown in black. **(f)** A subset of 100 randomly selected models ranked by their TM-score based on their similarity with WT Na_V_Sp1, and classified as Na_V_Sp1-like (black), Ms-like (magenta), horizontal CC (green), and splayed (cyan). Inserts show top-ranked representative models for each category. Most models (67%) fall above the cut-off. **(g)** Violin plots of the TM-align score distribution for all models (821 sequences), *Alphaproteobacter* class sequences (337 sequences), *Rhodobacter* order sequences (222 sequences), and *Roseobacter* family sequences (140 sequences) that include Na_V_Sp1. **(h)** Ranking of all 821 models based on their TM-align score showing the distribution of the RMSD values between each model and Na_V_Sp1.

Having established that AlphaFold predictions of the CTD of Na_V_Sp1 variants represent states that agree with the experimental data, we built AlphaFold2 multimer models of tetramers of 821 BacNa_V_ sequences out of the 825 sequences that we had obtained from the NCBI protein database. We excluded four sequences from this analysis, one because it lacked a voltage-sensing domain, two because AlphaFold failed to generate refined models, and one because it had annotation issues (**Dataset S05**). Despite the low sequence conservation along the CTD, most models with CTD sequences at least 28 amino acids long were predicted to contain a coiled coil in the distal CTD. Models with CTD sequences that were 22, 17, and 9 amino acids long had splayed C-terminal helices, while CTDs with 7 and 11 amino acids did not fold as helices in the predicted models (**Dataset S05**). Within the 821 models, six are from thermophiles and eight are psychrophiles (**Dataset S05**), for which the mean CTD length is 52.8 and 41.6 amino acids, respectively. We observed that in the top-ranked thermophile organism channel models the proximal CTD had a straight helical bundle conformation like that of Na_V_Sp1, while most psychrophile organism channel models had a kink in the CTD similar to the open state of Na_V_Ms (19) (**Fig. 6e**). We reasoned that channels predicted to have a CTD with a similar structure as Na_V_Sp1 could be susceptible to modulation by their CTD through a mechanism similar to what we described for Na_V_Sp1. To establish a metric to identify models with a CTD structure similar to that of Na_V_Sp1, we inspected 100 randomly selected models individually and classified these into four categories based on the structure of their proximal CTD as Sp1-like (black in **Fig. 6f**, straight helical bundle conformation), splayed (blue), Ms-like (magenta, kinked conformation), or horizontal (green, uncertain conformation). After ranking these 100 models by their structural similarity to Na_V_Sp1 using the TM-score (39) **(Fig. 6f)**, we found that most models classified as Sp1-like had a TM-score higher than 0.8562 while most models in the other categories had TM- scores less than this **(Fig. 6f)**. These findings indicate that models with a TM-score higher than 0.8562 are likely to have a CTD that is structurally similar to Na_V_Sp1. We observed that TM-scores trended toward larger values in models from organisms that are taxonomically closer to *Ruegeria pomeroyi* **(Fig. 6g)**, from which Na_V_Sp1 was cloned, suggesting that the structural similarity between models is at least partly associated with the evolutionary relations between bacterial species. Based on these observations, we ranked all 821 models based on their TM-score and found that more than half of them had a score higher than 0.8562 **(Fig. 6h)**, suggesting that the majority of these proteins have a fold similar to that of Na_V_Sp1, and could therefore be modulated by their CTD through a similar mechanism as we have described here. Importantly, the TM-scores that we used to rank the models were highly correlated with the RMSD between each of the models and Na_V_Sp1. The close agreement between both metrics indicates that our classification of the 821 models based on their structural similarity with Na_V_Sp1 is robust. This analysis of BacNa_V_ channel models constitutes an important resource for future interrogation of the potential biological role of these ion channels in diverse species of bacteria.

## Discussion

In this study, we show that the temperature-dependent activity of Na_V_Sp1 is intrinsic to its transmembrane domain, and we describe a mechanism that explains our experimental observations as well as those of others (17), whereby the cytoplasmic C-terminal domain of the channel tunes its activity in response to voltage and temperature. Based on our cross-species sequence analysis, AlphaFold predictions, molecular dynamics simulations, and experimental channel activity measurements, we conclude that in Na_V_Sp1 channels, voltage-dependent opening of the pore is associated with a conformational change in the helical bundle within the membrane-proximal portion of the CTD **(Fig. 7a)**. It is likely that this helical bundle in the proximal CTD experiences tension from both the pore-lining S6 helices on its N-terminus and the rigid coiled coil on its C-terminus, resulting in strong energetic coupling between the conformational state of these three regions. In WT Na_V_Sp1 channels, the membrane-proximal helical bundle adopts a conformation that is more rigid, which stabilizes the resting state of the channel and increases the energy barrier for pore opening **(Fig. 7a)**. A greater energy barrier for channel activation explains the increased temperature-dependent activity of WT Na_V_Sp1, while the stabilization of the closed state explains why stronger depolarization is required to fully activate these channels. Due to the strong energetic coupling between the coiled coil and the helical bundle in the proximal CTD, even small changes in the sequence of the coiled coil can alter the conformational landscape of the proximal CTD and increase its flexibility, with two major consequences for channel activity **(Fig. 7b)**: First, increased flexibility within the helical bundle in the proximal CTD destabilizes the resting state and lowers the energy barrier for pore opening, causing a hyperpolarizing shift in the voltage-dependence of activation and a decrease in temperature-sensitivity, respectively. This explains both the hyperpolarizing shift that we observed in most of the chimeras and the channels where the entire CTD or the coiled coil were truncated, as well as their decreased temperature dependence. Second, the increased flexibility enables the proximal CTD to adopt conformations in the open state where it can interact with the S4-S5 linker helix in the transmembrane domain, stabilizing the open state. The additional stabilization of the open state causes a further reduction in the temperature-dependent responses of the channel. This explains why the disruption of the interactions between the membrane-proximal CTD and the S4-S5 linker helix causes the temperature-dependence of the ΔCC construct lacking the coiled coil to increase **(Fig. 5f, g)**. Finally, we provide evidence that this mechanism could be conserved in a large portion of BacNa_V_ channels from other species of bacteria. Importantly, our finding that the intrinsic temperature-dependent activity of Na_V_Sp1 originates in its transmembrane domain, which has a fold that is similar to eukaryotic voltage-gated channels and is highly conserved across prokaryotic Na_V_ channels, raises the possibility that the temperature-dependence of eukaryotic channels, including those in neurons, as well as BacNa_V_ channels from other bacterial species, could arise from a mechanism similar to that in Na_V_Sp1. Although the molecular mechanism responsible for the intrinsic temperature-dependence of Na_V_Sp1 activity remains to be determined, it appears to be tightly associated with the gating of the pore rather than voltage-sensor activation (17). Consistent with this hypothesis, pore gating has been proposed to have elevated intrinsic temperature sensitivity in other ion channels with a similar transmembrane domain fold as BacNa_V_ channels, including voltage-activated K^+^ channels (40) and TRPV1 channels (41).

**Figure 7.**
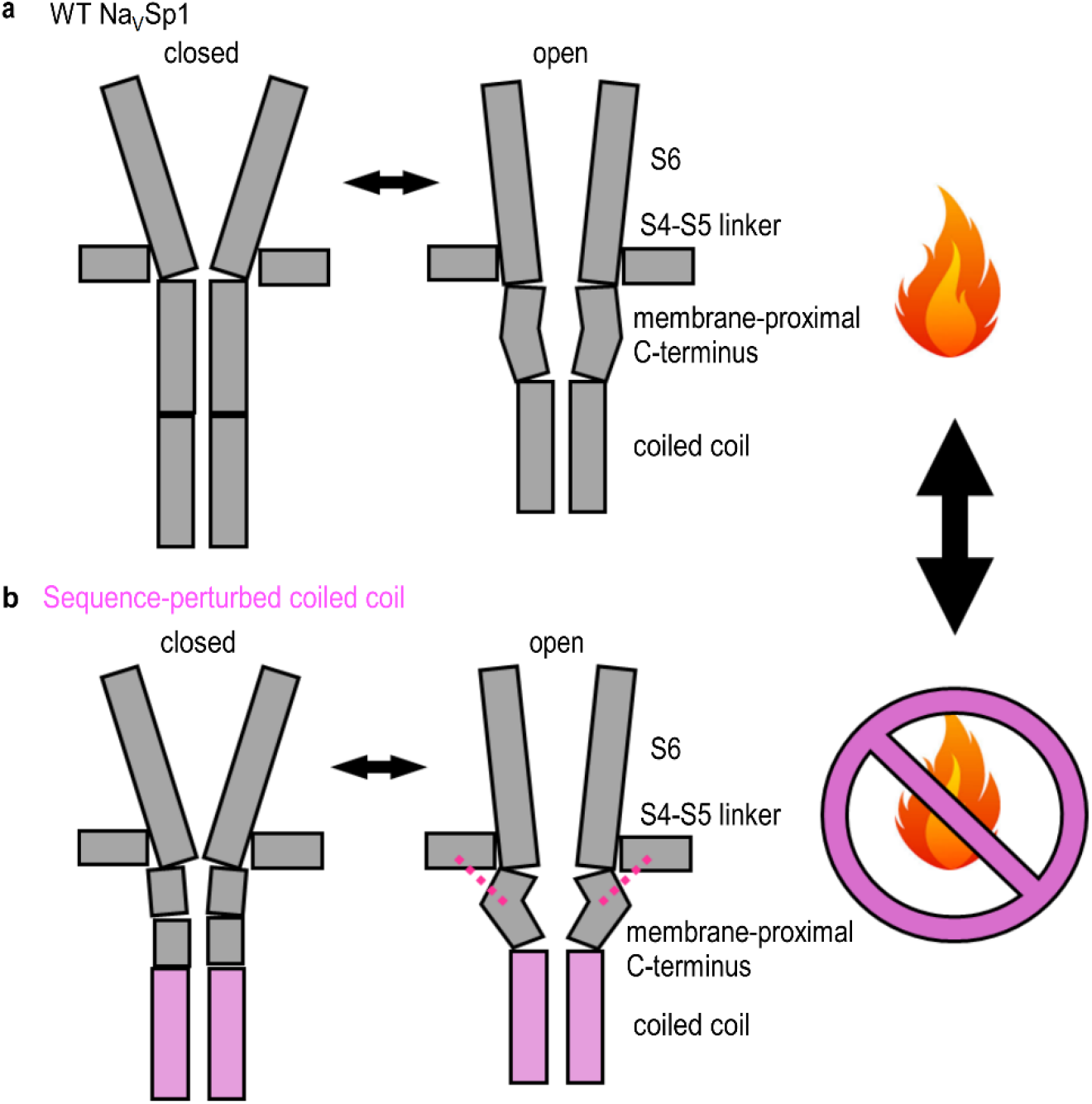
Mechanism of Na_V_Sp1 channel tuning by its C-terminal domain. **(a)** Cartoons representing the gating transition between closed and open states in the WT Na_V_Sp1. The conformational change in the membrane-proximal CTD tunes the voltage- and temperature-dependent response of the channel, resulting in strongly temperature-dependent channel activity. **(b)** Cartoons representing the gating transition between closed and open states in Na_V_Sp1 channels where the sequence and structure of the coiled coil (lilac) has been perturbed. The conformational change in the membrane-proximal CTD is associated with a lower energy barrier, and the CTD no longer contributes to stabilizing the closed state. In addition, proximal CTD can interact with the S4-S5 linker, stabilizing the open state, resulting in weak temperature-dependence.

The development of this mechanistic model explaining how the CTD of Na_V_Sp1 tunes its activity was born from the agreement between the structural features of the CTD predicted by AlphaFold and the experimentally determined behavior of multiple constructs with altered sequence. While AlphaFold is known to have limited accuracy for predicting the structural effects of mutations, it was found to make remarkably accurate predictions of coiled coil geometry (42). This accuracy arose from the structural embeddings in the AlphaFold model and not the sequence alignment (42). This may be due to the large number of distinct coiled coil structures available in the protein database (43), many of which have distinct sequences despite adopting a similar fold.

These unique properties of coiled coil sequences may contribute to the accuracy in the predictions that we observed. Importantly, we found that the AlphaFold predictions for each of the channels that we examined included models with open and closed pores, where the latter may represent closed or inactivated states. Regardless of whether the non-conducting conformations represent closed/resting or inactivated states, we only observed interactions between the S4-S5 linker and the CTD of the channel in models in which the pore was in an open state.

To examine the implications of the mechanism described above, we analyzed the predictions of a simple 7-state sequential kinetic model that was sufficient to qualitatively reproduce all key features of WT Na_V_Sp1 activity **(*SI Appendix,* Fig. S8a)** – the model showed strongly temperature-dependent responses **(*SI Appendix,* Fig. S8b, c)**, a depolarized voltage of activation that was only mildly affected by temperature **(*SI Appendix,* Fig. S8d, e)**, and current kinetics that were similar to the experimental recordings **(*SI Appendix,* Fig. S8c, e)**. Although in this model the temperature-dependence of channel activation is directly associated with the opening rate constant k_2_, more complex models with additional temperature-dependent transitions between voltage-sensor activation and pore opening should exhibit similar behaviors. We also note that the use of equilibrium thermodynamics is justified, as we found the temperature responses of Na_V_Sp1 to be reversible **(*SI Appendix,* Fig. S8f)**. To represent the effect of the truncation of the coiled coil (ΔCC), we incorporated into the model an additional open state O* representing the formation of the interactions between the proximal CTD and the S4-S5 linker, and modified the rate constants of activation and inactivation to reflect a destabilization of the closed state and a stabilization of the open state. This model succeeded in representing all key features associated with the activity of ΔCC **(*SI Appendix,* Fig. S8b-e)** – weakly temperature-dependent responses **(*SI Appendix,* Fig. S8b, c)**, a left-shifted voltage of activation relative to WT **(*SI Appendix,* Fig. S8d, e)**, and slower kinetics of activation and inactivation **(*SI Appendix,* Fig. S8c, e)**. Together, these results lend further support to the mechanism that we proposed.

Consistent with the mechanistic interpretation of our results, amino acid sequence changes in the CTD of the Na_V_Sp1 channel were shown to be energetically coupled to the pore gate and not the voltage-sensor domain (17). This mechanism also explains previous observations that the introduction of glycine substitutions or other flexibility-increasing mutations into the proximal CTD cause both a hyperpolarizing shift in the voltage-dependence of activation and a reduction in temperature sensitivity (17, 18): these alterations destabilize the resting state, lower the energy barrier for pore opening, and allow the proximal CTD to establish interactions with the S4-S5 linker that stabilize the open state.

The mechanism we described is also consistent with experimental observations in other BacNa_V_ channels, which show how variations in the structural features of the CTD can impact channel activity; In the Na_V_Ae1 channel from the haloalkaliphilic gamma proteobacterium

*Alkalilimnicola ehrlichei*, the membrane-proximal CTD appears to be particularly stable because it was resolved in structures of the isolated closed pore, where it folds as a rigid helical bundle (17, 18) (***SI Appendix,* Fig. S1**). Interestingly, the Na_V_Ae1 channel exhibits no detectable voltage-dependent activity in heterologous expression systems unless mutations are introduced that destabilize the membrane-proximal CTD (17, 18), suggesting that in these channels this region is too rigid to allow the pore to open at room temperature. Furthermore, deletion of the CTD of the Na_V_Ab channel from *Arcobacter butzleri* stabilized the open state, allowing the determination of an open structure by X-ray crystallography (22). These observations establish that increased rigidity of the membrane-proximal CTD can interfere or prevent gating in some BacNa_V_ channels. According to AlphaFold predictions, the CTD in the top-ranked models of BacNa_V_s from six thermophilic organisms had a straight helical bundle fold, similar to Na_V_Ae1. A rigid C-terminal fold would oppose the conformational changes required for the pore to open, and thus help stabilize the closed state at higher temperatures that could reduce the stability of the proximal CTD. In contrast, the membrane-proximal CTD in the Na_V_Ms channel from the alpha proteobacterium *Magnetococcus marinus* has the opposite properties – full-length open state structures of this channel show one half of the membrane-proximal CTD as an α-helical extension of the S6 helices and the other half adopts a random coil configuration (19, 21) **(*SI Appendix,* Fig. S1)**. Unlike Na_V_Ae1, deletion of the coiled coil of Na_V_Ms does not alter its voltage-dependence of activation, instead affecting channel expression as well as the kinetics of inactivation and recovery from inactivation (19), indicating that the stability of the membrane-proximal CTD does not make an energetic contribution to pore gating in this channel. This is consistent with DEER measurements indicating extensive flexibility within the membrane-proximal CTD of Na_V_Ms (21). Importantly, the helical portion of the membrane-proximal CTD of Na_V_Ms is shown to interact with the S4-S5 linker (19, 35, 36) as we described for Na_V_Sp1. These interactions in Na_V_Ms were shown to stabilize the open state and modify the kinetics of activation and inactivation, consistent with the mechanistic interpretation of our results. The CTD in most top-ranked models of channels from psychrophilic organisms had a structure similar to that of Na_V_Ms, which would be predicted to stabilize the open state, which we propose could be important to enable channel activity at low temperatures.

The biological role of the CTD in Na_V_Sp1 or other BacNa_V_ channels remains to be determined. Because the resting membrane potential in bacteria is too negative (44, 45) for channels to activate based on the voltage-activation relations of the few BacNa_V_ channels whose activity has been studied, specific regulatory mechanisms or factors must exist in bacteria to enable channel activity (46). The strong temperature- and pH-sensitivity found in three channels from alkaliphilic *Bacilleae* bacteria was thus proposed to enable their activation at the very negative resting membrane potentials, allowing channels to contribute to motility, chemotaxis, and pH regulation (6, 46). Due to the small size of a bacterial cell, even brief openings of a few channels could have a significant impact on the membrane potential and the concentrations of ions inside the bacteria (47). Our analysis of the predicted structural features of over 800 non-redundant BacNa_V_ channel sequences suggests that in more than half of these sequences, the CTD adopts a structure that is similar to that of Na_V_Sp1. It is therefore likely that the voltage-dependence of these channels may be susceptible to regulation through a similar mechanism as in Na_V_Sp1, where the conformational landscape of the membrane-proximal helical bundle, as well as the interactions it can form with the S4-S5 linker in the open state, determine the voltage of channel activation and its temperature-dependence. The strong energetic coupling that we detected between the structure of the coiled coil and the conformational landscape of the helical bundle in the membrane-proximal CTD raises the intriguing possibility that environmental or cellular factors such as acidity, ionic strength, or osmolarity, could influence the quaternary structure of the coiled coil, and thus of the entire CTD, to enable or disable temperature-dependent channel responses. Our work thus identifies a regulatory mechanism that can help to elucidate the enigmatic biological function of voltage-gated Na^+^ channels in diverse bacteria.

## Materials and Methods

### BacNa_V_ channel amino acid sequences and taxonomy

Our bioinformatics analysis was performed by searching the term “voltage gated sodium channel” in the National Center for Biotechnology Information (NCBI) Protein Database (https://www.ncbi.nlm.nih.gov/protein/). Results were further narrowed by limiting the search to entries that contained a range of 100 to 500 residues. To eliminate redundancy of sequences we utilized CDHIT software (48), allowing us to remove amino acid sequences to produce a set with no more than 90% identity between any two members, and reducing our original 1,392 entries to 825 **(Dataset S01)**. Out of the 825 entries, we were able to extract taxonomic associations for all proteins using the NCBI database and obtained environmental associations for 432 entries using BacDive (49), a bacterial diversity metadatabase that details taxonomic and strain information for large number of bacterial species identified in metagenomic samples from soil, water, animal, or plant origin. Notably, the 432 entries on this list represent 392 unique bacterial species listed on **Dataset S02**. This discrepancy arises because a small number of bacteria had more than one Na_V_ channel sequence. Our taxonomic data were compared to a previously constructed bacterial phylogenetic tree. Additionally, using BacDive records of species’ global distribution based on their 16S sequences, we classified bacterial habitats as predominantly aquatic, soil, animal, or plant if over 60% of the total BacDive sample counts where a bacterial species was identified were associated with the same type of environment. Otherwise, species were labeled as multi- environment.

The phylogenetic tree was created using data from the GTDB (50) and NCBI taxonomy databases, based on bacterial classes filtered through top GTDB entry counts and BacNa_V_-containing species. Briefly, we scanned the GTDB taxonomic classification, sorted out all the families, genus, classes, etc. and their counts established in the database. We then scanned our taxonomic data to find entries with 10 or more BacNavs for every taxonomic classification. Then we determined which GDTB taxonomic classifications overlapped with our taxonomic data for the species containing Na_V_Sp1 channels. The rationale for this is that some taxonomic classifications found in the NCBI were not found in the GDTB. We found that class was the best taxonomic classification since itincluded all BacNa_V_ entries. If any of the classes derived from our NCBI BacNav sequence list had a minimum of 10 distinct channel entries and did not appear in a list of the top classes with the most entries found in the GTDB, we replaced the lowest GTDB entry with the class from the BacNav sequence set. The 12 classes displayed on the tree are not a comprehensive listing of all known classes of bacteria.

### Molecular biology

Mammalian expression plasmids for HA-tagged Na_v_Sp1 full-length – IRES2 – GFP (# 128087) and Na_v_Sp1 ΔCC – IRES2 – GFP (# 128084) were purchased from Addgene. Additional coiled coil deletions and chimeras were generated by PCR and restriction-ligation cloning using full-length Na_v_Sp1 as template. Briefly, we used a long mutagenic reverse primer (Integrated DNA Technologies) and a short forward primer (Integrated DNA Technologies Ultramer), that contained the stop codon for the channel and an EcoRI restriction site immediately downstream. Long primers were heated at 98 °C for 10 minutes to improve PCR yield. The PCR product was digested with EcoRI enzyme, self-ligated using T4 ligase, and transformed into E. cloni 10G chemically competent cells (Biosearch Technologies). All primer sequences are included in **Dataset S03**. Coiled coil chimeras were generated from the BacNa_v_ channel sequences from *Bacillus halodurans* (Na_v_Bh; protein -TES55043.1; DNA - SNUY01000019.1)*, Magnetococcus marinus* (Na_v_Ms; protein - WP_011712479.1; DNA - NC_008576.1)*, Pseudooceanicola batsensis* (Na_v_Ob; protein - WP_009804693.1)*, Alcanivorax borkumensis* (Na_v_Ab1; protein - WP_062810689.1; DNA - NZ_BDAT01000007.1), and *Rhodobacteraceae bacterium KLH11* (Na_v_Rbl; protein - EEE38440.1). We defined the coiled coil region of the Na_v_Rbl and Na_v_Ob channels, for which no structural information is available, using a multiple sequence alignment in combination with the predicted coiled coil probability (51). We used the protein sequence to generate a codon-optimized DNA sequence for cloning and expression. For the Na_v_Sp1-Ab1cc ΔR248 and Na_v_Sp1-Mscc ΔL248 deletion chimeras, as well as the cpHALO-Na_v_Sp1 L253C + G254C construct, the entire coding region was purchased as a double-stranded DNA fragment (TwistBio) and cloned into the expression plasmid using NheI and EcoRI restriction enzymes. Fragment sequences are included in **Dataset S03**. The cpHALO Tag sequence that we used matches that in Addgene plasmid #138325 and included the C-terminal linker sequence Ser-Gly-Leu-Arg-Ser-Ala-Thr-Ala (52). The cpHALO-Na_v_Sp1 WT construct was obtained by restriction-ligation cloning of the WT coiled coil sequence insert using BsmBI and HindIII enzymes. All constructs were verified by Sanger sequencing.

### Cell culture

We kept human embryonic kidney cells (HEK293, ATCC CRL-1573) at 5 % CO_2_ and 37°C. We grew cells in Dulbecco’s modified Eagle’s medium (DMEM) with high glucose, pyruvate, L-glutamine, and phenol red, supplemented with 10 % heat-inactivated fetal bovine serum (v/v) and 10 mg/mL gentamicin. Before transfection, we detached cells by incubation with 0.25 % Trypsin, resuspended them in DMEM, and seeded cells on 20 x 20 mm No. 1 glass coverslips in 35 mm dishes. We transfected cells immediately after seeding using FuGENE HD (Roche Applied Science) following manufacturer’s instructions. We conducted patch clamp recordings 18 - 48 hours after transfection.

### Patch clamp electrophysiology

We carried out all experiments using the whole-cell configuration of the patch-clamp technique. Before experiments, we broke the 20 x 20 mm glass coverslips that contained cells into fragments using metal tweezers, transferred one of the fragments into a 35 mm dish containing extracellular solution, and broke it into smaller fragments that we used for recording. We obtained a single experiment per coverslip fragment. After gigaseal formation and breaking into whole-cell configuration, we lifted cells a few millimeters from the coverslip before starting the experiments. We acquired and digitized data at 5 kHz using a dPatch amplifier system and SutterPatch software (Sutter Instruments) under voltage clamp. We pulled pipettes from borosilicate glass (1.5 mm O.D. × 0.86 mm I.D. × 75 mm L; Harvard Apparatus) using a Sutter P-97 puller and heat-polished to final resistances between 0.2 – 2 MΩ using a MF-200 microforge (World Precision Instruments). We used an agar bridge (1 M KCl; 4% w/v agar; Teflon tubing) to connect the ground electrode chamber and the main recording chamber. We performed all recordings using a temperature-controlled Delta T imaging chamber (Bioptechs). The Delta T chamber enables uniform and rapid temperature changes by delivering heat directly from the cover glass, which has an electrically conductive coating on the underside opposite where cells sit. We recorded temperature in all experiments using a small thermistor probe placed a few millimeters away from the patch pipette tip. We measured current-voltage relations at various temperatures by changing the temperature setpoint on the chamber, and current-temperature relations using the heat shock mode that results in more rapid changes in temperature. For rapid cooling we flowed ice water through a stainless-steel cooling ring (Bioptechs) that we positioned in the recording chamber before starting the experiment. For experiments requiring perfusion of dithiothreitol (DTT)-containing solution, we used a gravity-fed motorized RSC-200 perfusion system (BioLogic).

The intracellular solution was composed of 150 mM CsMethanesulfonate, 8 mM NaCl, 20 mM HEPES, 10 mM EGTA, 2 mM ATP (pH 7.5). The extracellular solution consisted of 155 mM NaCl, 1 mM CaCl_2_, 1 mM MgCl_2_, 5 mM KCl, 20 mM HEPES, 10 mM glucose (pH 7.4). For experiments we used a 30 % (w/w) (9.8 M) H_2_O_2_ solution with stabilizer as stock and a 1 M stock of DTT prepared in water. We used a holding potential of −90 mV, a pulse duration of 300 ms, and an inter-pulse interval of 3 seconds for all experiments. We measured current-voltage relations by stepping voltage from −90 to 60 mV in increments of 10 mV. For current-temperature relations, voltage was repeatedly stepped between −90 and 30 mV. A dotted red line denotes the zero-current level in all figures depicting current traces.

We analyzed data using Igor Pro 9 (Wavemetrics) and Microsoft Excel. We obtained current-voltage (I-V) and current-temperature (I-T) relations by quantifying the mean peak current for each voltage pulse. Individual I-T relations were normalized to the mean peak current at room temperature (20 – 22 °C). Mean ± SEM I-T relations were calculated by averaging the mean peak current across experiments from a single pulse per experiment per 2 °C interval (41). We obtained Q_10_ values from log(I)-T relations by fitting a line to the steepest region of the curve encompassing an interval > 5 °C and involving at least three datapoints and substituting the resulting slope (*m*) into Eq. 1:

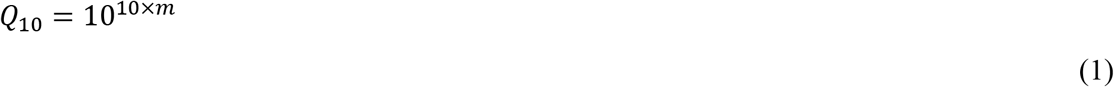

To obtain conductance-voltage relations, current-voltage relations were first fit using Eq. 2:

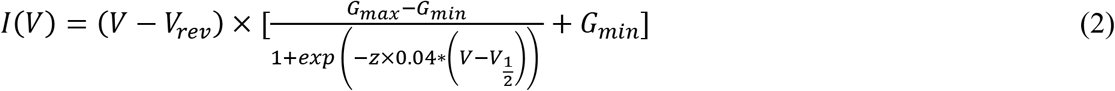

We then calculated the conductance by Eq. 3:

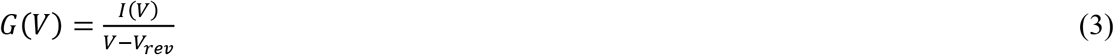

We obtained the voltage of half-maximal activation (*V_1/2_*) by fitting the *G*-*V* relations to the Boltzmann equation, given by Eq. 4:

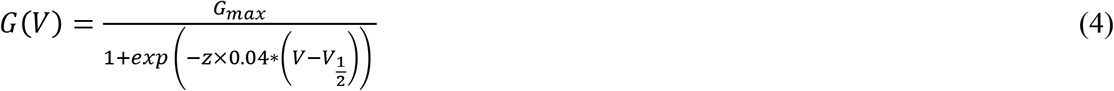

The time constants of channel activation and inactivation were obtained from mono-exponential function fits to current traces obtained from experiments to determine the current temperature relations.

### Mathematical modeling of channel activity

We solved the system of differential equations corresponding to the kinetic model described in ***SI Appendix,* Fig. S8a** with the Backwards Differentiation Formula method implemented on Igor Pro 9. Ionic currents were calculated according to Eq. 5,

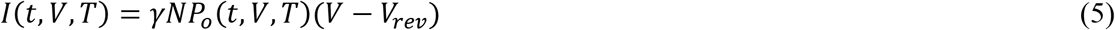

The fit parameters were manually optimized by visual assessment of the agreement between calculated and experimental data. The voltage-dependent rate constants (*k*_0_, *k*_1_) had a form given by Eq. 6:

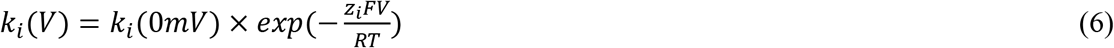

In the above, *k_i_*(0*mV*) is a constant, *z_i_* is the valence associated with the transition, *F* is Faraday’s constant, *R* is the gas constant and *T* is the temperature. The temperature- and voltage-dependent rate constant *k*_2_ is given by Eq. 7:

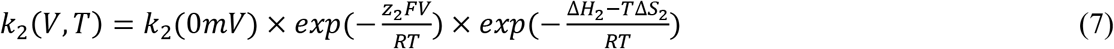

In Eq. 7, *ΔH*_2_ and *ΔS*_2_ are the changes in enthalpy and entropy associated with reaching the transition state associated with that rate constant. The temperature-dependent rate constant *k_3_* is given by Eq. 8:

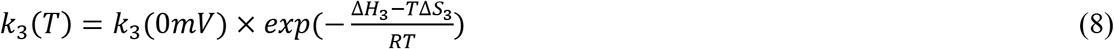

Importantly, *ΔH*_3_ and *ΔS*_3_ were negligible, so that *k*_2_ was the only effective contributor to the temperature dependence. The *I*-*V*, *I*-*T*, and *G*-*V* relations arising from the model were obtained from the predicted currents as described for the experimental data. We disregarded recovery from inactivation by assigning it a very slow rate constant, because our voltage step protocol allowed for channels to fully recover from inactivation after each step. Therefore, we set full occupancy of the first closed state *C*_0_ as the initial condition for each simulated voltage step.

### Cysteine crosslinking, SDS-PAGE, and in-gel fluorescence

We seeded cells at 50-70% confluency on 6-well plates that were pre-treated for 30 minutes with bovine Fibronectin (0.1 mg/mL in PBS) and transfected as previously described. After 12 hours, we washed wells with extracellular recording solution, followed by a 30-minute incubation at 37 °C with JF549 dye (Promega) prepared in FluoroBrite DMEM at a concentration of 200 nM from a DMSO stock solution at 200 μM. After washing the dye off, we incubated cells at room temperature for 5 minutes in extracellular solution with or without 10 or 40 mM H_2_O_2_. We washed wells one more time, followed by a 5-minute incubation in extracellular solution with or without 20 mM DTT. After washing once, we detached cells by pipetting, spun cells down at 1,000 G for 2 minutes, lysed cells (Pierce Surface Protein Biotinylation and Isolation Kit lysis buffer, Thermo Scientific), and spun lysate down at 10,000 G for 2 minutes. The supernatant for each sample in each of three independent replicates was run under denaturing and non-reducing conditions using NuPAGE 4-12% Bis-Tris gels (Invitrogen) and NuPAGE MOPS SDS running buffer. Gel fluorescence was imaged in a Typhoon RGB (Amersham) or a ChemiDoc (BioRad) gel imager. We quantified band fluorescence intensities using FIJI (53), and normalized the fluorescence intensity for the monomer, dimer, trimer, or tetramer to the sum of the fluorescence intensities from the four species.

### Molecular dynamics simulations and AlphaFold2-Multimer modeling

Structure prediction was performed with a local installation of AlphaFold2-multimer (2.2.0) (54). We performed molecular dynamics simulations of open and closed states selected from pools of AlphaFold 2 (AF2) model predictions for WT NavSp1, ΔCC-2+DQRL, ΔCC-2+DQR, the Ms coiled coil chimera, and its single-residue variant Ms ΔL. All simulations were set up using CHARMM-GUI (55–57) and simulations were performed with the GROMACS 2022 software package (58). The Charmm36m force-field with the TIP3P water model was used (59). The models were embedded in a DPPC membrane for simulation with 150 mM NaCl plus neutralizing ions. The simulation systems each contained around 181,000 atoms: 132,000 atoms of water and ions, 16,000 protein atoms, and 32,000 atoms in the lipid membrane. Simulation box sizes for each simulation are approximately 13.4nm x 13.4nm x 17.8nm. For each simulation setup 1 µs of simulation was run with two subsequent replicas of 500ns each, unless otherwise noted. Each system was energy minimized using steepest descent followed by a multi-step equilibration lasting a total of 375 ps. In the first two steps, the system was equilibrated in a canonical (NVT) ensemble with an integration time step of 1 fs for 50 ps each, maintaining a temperature of 323.15 K using the Berendsen thermostat (60). In the case of the higher temperature ΔCC-2+DQR simulations, these were run at 363.15 K and with only two replicates of 500ns each. During this multi-step equilibration, position restraints were applied on lipid and protein heavy atoms, starting at 1,000 and 4,000 kJ/(mol/nm2), respectively, and then gradually lowered in every step until reaching zero in the final step. During the equilibration steps, we established a pressure of 1 bar using the Berendsen barostat with a characteristic time of 1 ps(60). During the simulations, no restraints were applied on the systems. We applied the velocity rescale thermostat with a characteristic time of 1 ps to keep a constant temperature of 323.15 (61). We applied the Parrinello-Rahman barostat with semiisotropic pressure coupling and a characteristic time of 5 ps to keep a constant pressure of 1 bar (62). Thermostats were applied independently on protein, membrane, and solvent. Throughout all simulations, non-bonded interactions were cut off at 1.2 nm. Optimal usage of computational resources for these simulations was assessed using MD Benchmark (63). Simulations and models were analyzed in Python using MDTraj (63), TMalign (39), and Hole (64) and figures were made using PyMOL (http://www.pymol.org).

### Rosetta coiled coil design

Using the RosettaScripts scripting language (64), we developed a protocol for *de novo* design of an artificial C4-symmetric coiled coil domain that could be fused to a BacNa_V_. Our protocol used Rosetta’s parametric design tools (65) to sample coiled coil conformations, varying bundle radius (*r*_0_), bundle twist (*ω*_0_), and helix roll (Δ*ω*_1_). It then built C4 symmetry and applied Rosetta’s FastDesign tool (66) to design the sequence. Design was constrained with amino acid composition constraints (67) to favor a hydrophobic core and polar surface and to limit aromatic, alanine, and methionine residues, as well as with net charge constraints (68) to promote a mildly negative net charge to minimize coiled coil toxicity on expression. Additional guidance terms were applied to minimize interfacial voids, minimize buried unsatisfied hydrogen bonding groups, and maximize hydrogen bond networks (69)(68). Following design, the protocol relaxed the structure using the RosettaQM bridge (manuscript in preparation) calling out to GAMESS (70), which applied the DFTB semi-empirical quantum chemistry (QC) method with the fragment molecular orbital (FMO) approximation (71). Finally, the protocol evaluated seven metrics: the number of buried unsatisfied hydrogen bond donors and acceptors, the shape complementarity between helices, the interface cavity volume, the total energy of the coiled coil computed with Rosetta’s ref2015 force field (72), the interface energy computed with ref2015, the interface energy computed with QC (DFTB/FMO) by RosettaQM/GAMESS, and the QC energy with Rosetta’s solvation terms (fa_sol, fa_intra_sol_xover4, and lk_ball_wtd) added. We generated 1,683 coiled coil designs and manually inspected the 165 designs that were Pareto optimal based on the seven metrics calculated. Of these, we chose two that showed good core packing, few interfacial holes, favorable hydrogen bond networks, and helical twist resembling that of the natural BacNa_V_ coiled coil. The sequences of the designs that we ranked first and second were LEHDIYQHLNNIERYLRRIE and EQNDLYRRIQQIEHYLRKLQ, respectively. See the data availability section below for links to the RosettaScripts scripts used to engineer the artificial coiled coil.

### Coiled coil geometry analysis

The geometry of the coiled coils in the top-ranked models for each of the chimeras used in this study was analyzed using the SamCC Turbo (34) webserver (https://lbs.cent.uw.edu.pl/samcc_turbo). The pdb file of each tetrameric complex model was uploaded and analysis was done using the Auto mode to identify the location of the coiled coil. The coiled coil start was set at amino acid position 236 for WT and all chimeras, and the last amino acid in each model was defined as the end of the coiled coil.

## Data and materials availability

Sequence information for each of the 825 BacNa_V_ channels analyzed in this study is provided on the **Dataset S01** file. Information about the environments associated with some of the bacterial species that contain BacNa_V_ channels is included in the **Dataset S02** file. A list of PDB ID numbers for structures of BacNa_V_ channels together with references is provided in the **Dataset S03** file. Primers and double stranded DNA fragment sequences used in cloning are included in the **Dataset S04** file. Sequences with particular characteristics are listed in the **Dataset S05** file. All scripts used in this study, as well as all AlphaFold models and MD simulation trajectories included in this study, are freely accessible at: http://www.github.com/flatironinstitute/BacNav-Cterm-regulation/Correspondence and requests for materials should be addressed to: Andrés Jara-Oseguera Sonya M. Hanson

## Acknowledgements and funding sources

We thank Despoina Mavridou, Jeffrey E. Barrick, Mary Ann Moran, Christa B. Smith, Eric Senning, Marcel Goldschen-Ohm, Susanne Ressl, and Sören von Bülow for helpful discussions. We thank Bargeen Turzo and Doug Renfrew for help with high-throughput application of AlphaFold2-Multimer on the Flatiron Institute clusters. We thank Helen Chen and Averi Pike for technical assistance, Jason McLellan, Eric Senning, and Susanne Ressl for sharing equipment and reagents. We would also like to thank Martin Centola for help with experiments in earlier stages of this work that did not make it to the final paper. SMH was supported by a Humboldt Research Fellowship for Experienced Researchers from the Alexander von Humboldt Foundation during early stages of this research. This research was supported by startup funds from the University of Texas at Austin (to AJO). The Flatiron Institute is a division of the Simons Foundation.

## Contributions

S.M.H. and A.J.O. conceptualized, supervised, and administered the project. S.M.H., A.J.O. and G.H. acquired funds. C.G. and A.J.O. performed electrophysiological experiments. A.Z.C., A.V., and A.J.O. carried out molecular cloning. A.J.O. performed in-gel fluorescence experiments and mathematical modeling of channel activity. A.Z.C. performed amino acid sequence data mining and curation. S.M.H. carried out molecular modeling, molecular dynamics simulations, and bioinformatics analysis. V.M. carried out Rosetta modeling. A.Z.C., C.G., S.M.H., and A.J.O. analyzed and curated data, and contributed to visualization. A.J.O. wrote the original draft of the paper. A.Z.C., V.M., G.H., S.M.H., and A.J.O. reviewed and edited the paper.

