## Supplementary Appendix for "The mechanism of regulation of prokaryotic Na^+^ channels by their C-terminal domain"

**This PDF includes:**

Figures S1 to S8.

**Other supporting materials for this manuscript include the following:**

Datasets S01 to S05.

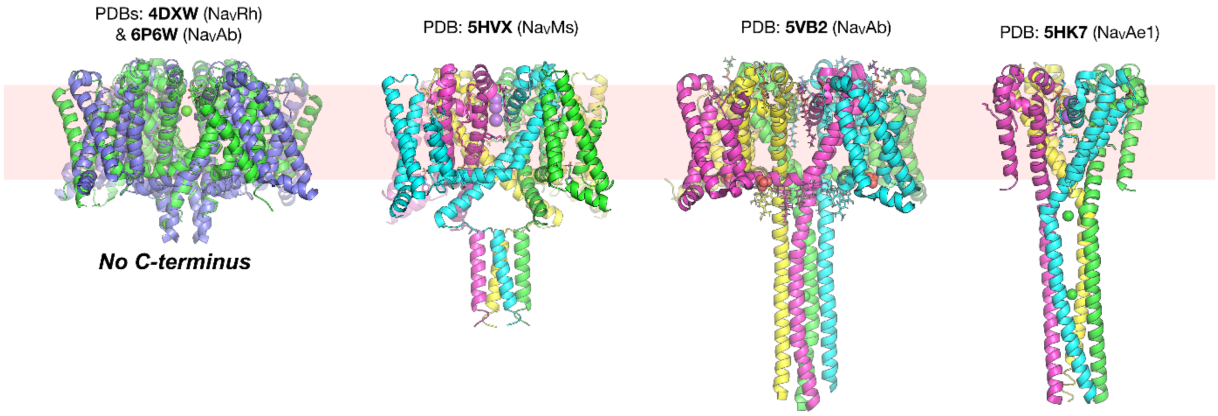


**Figure S1. Experimentally determined structures representative of BaNa$_V$ channels.** Most structures of BacNavs (see **Dataset S03**) do not contain C-termini, exemplified by the leftmost structures of NavRh (72) and NavAb (73) in purple and green. Exceptions to this are structures with resolved C-termini, from left to right: NavMs (19), NavAb (22), and NavAe1 (17). The right-most structure is of the pore-only construct, referred to as NavAe1p, which contains no voltage sensing domains and includes only the pore domain and CTD.

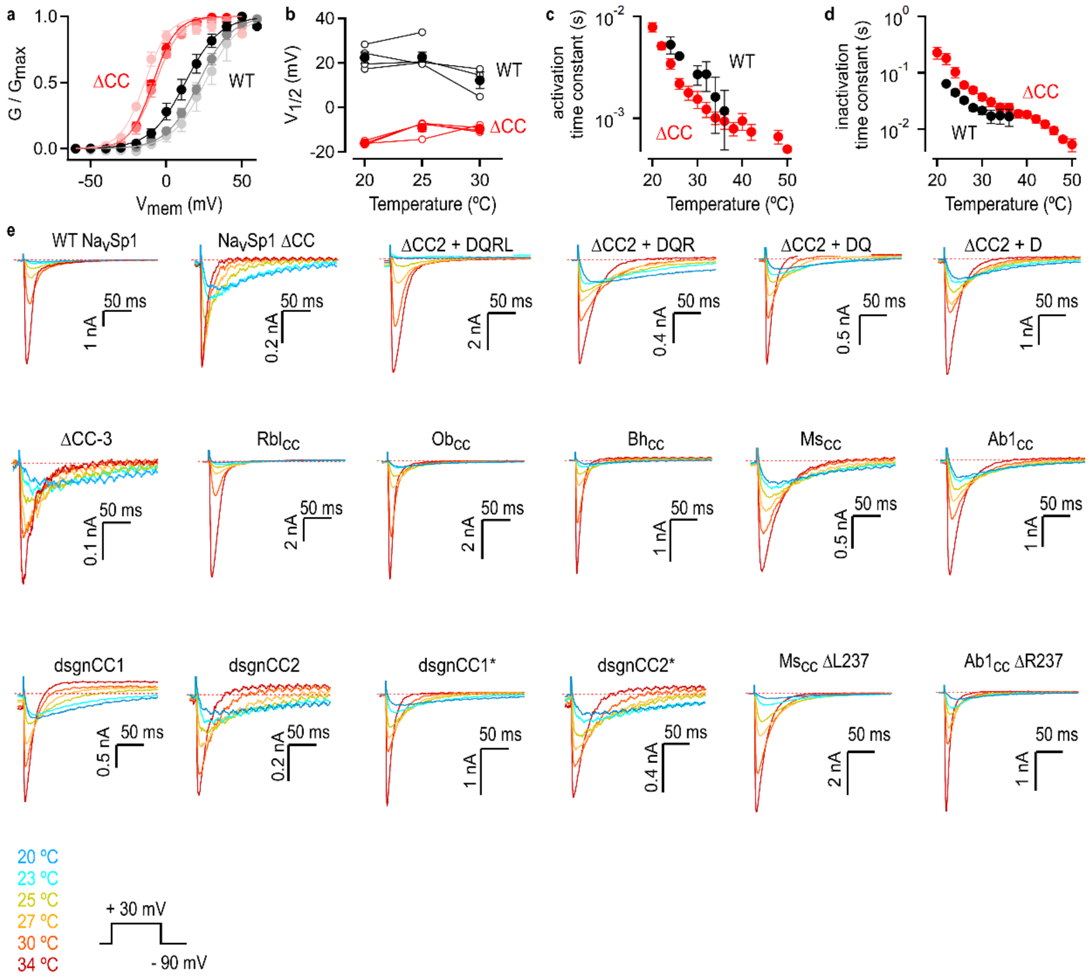


**Figure S2. Quantifying the temperature- and voltage-dependence of BacNa$_v$ channel constructs. (a)** Conductance-voltage relations at three temperatures obtained from experiments as in Fig.1d (mean ± SEM; WT, n = 3 – 6; ΔCC, n = 4). Continuous curves are fits to the Boltzmann equation. **(b)** $V_{1/2}$ values obtained from conductance-voltage relations obtained at different temperatures as in (a). Data from individual experiments are shown as circles and lines between data points that were obtained from the same experiment at distinct temperatures. The mean ± SEM is shown in squares. **(c and d)** Time constants of channel activation (c) and inactivation (d) as a function of temperature, obtained from experiments as in Fig. 1g (mean ± SEM, WT, n = 8; ΔCC, n = 7). **(e)** Representative current traces elicited during a temperature ramp by repetitive voltage steps from -90 to 30 mV in cells expressing Na$_V$Sp1 channel constructs. Current traces for only six temperatures are shown for visual clarity.

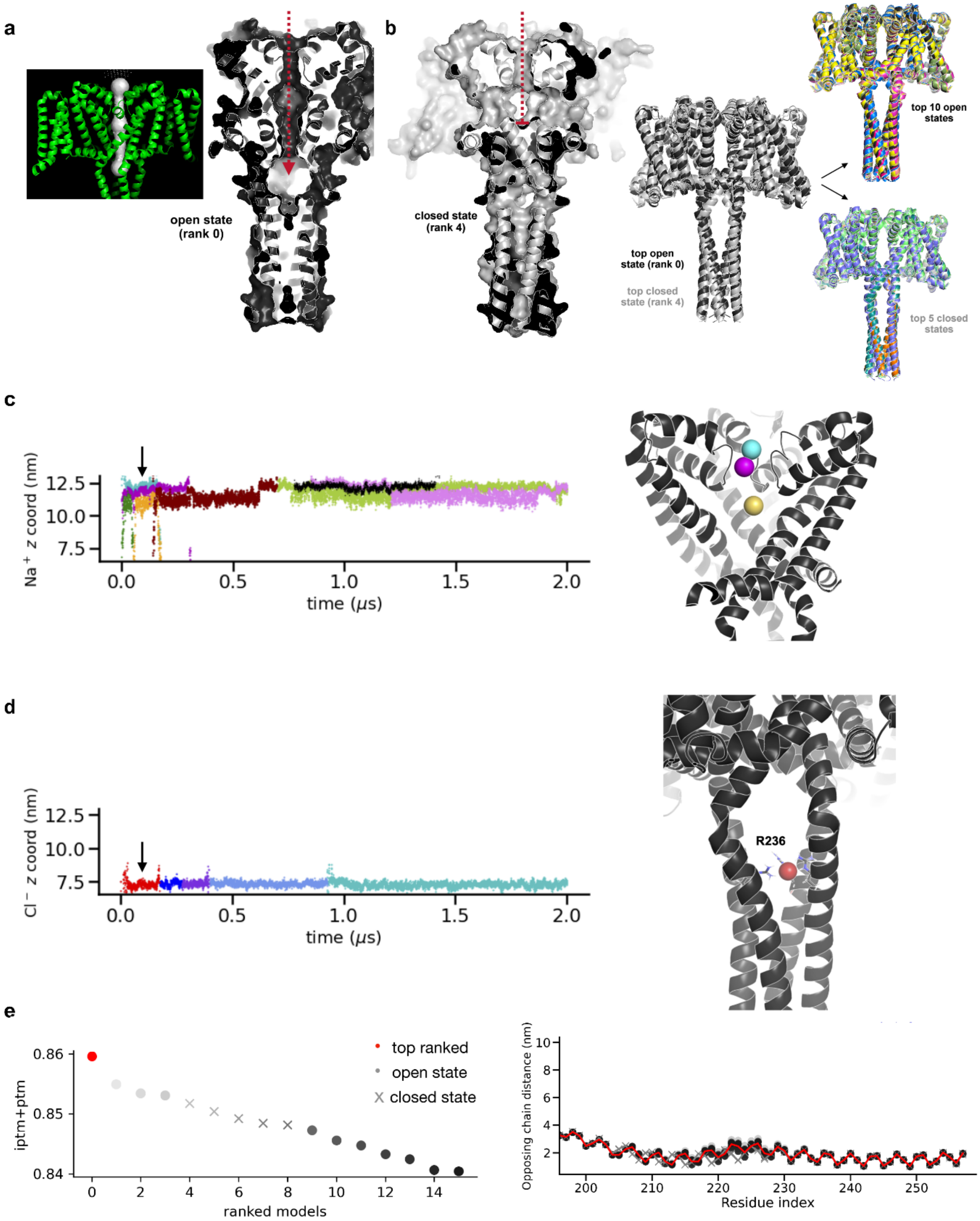
a
open state
(rank 0)
b
closed state
(rank 4)
top open
state (rank 0)
top closed
state (rank 4)
top 10 open
states
top 5 closed
states
c
Na$^+$ z coord (nm)
time (μs)
d
Cl$^-$ z coord (nm)
time (μs)
R236
e
iptm+ptm
ranked models
top ranked
open state
closed state
Opposing chain distance (nm)
Residue index

**Figure S3. AF2 models of WT Na$_V$Sp1 sample both open and closed states. (a, left)** Analysis of the top model of WT NavSp1 with Hole (74) shows it to be in an open state. **(a, right)** Surface representation shows an open pathway within the pore domain for the top-ranked NavSp1 model. **(b, left)** The closed pore models of NavSp1 as predicted by AF2 (grey), showing an occlusion of the inner pore formed by the bottom portion of the S6 helices. **(b, right)** Overlays of the top open and closed state models. **(c, left)** Trajectories for each of the observed $Na^+$ ion permeation events during the simulation started from the top-ranked NavSp1 model. Each event in which separate $Na^+$ ions traverse the pore is shown in a different color. Note that most ions leave the pore from the same side as they entered. Here the only ion that crossed completely from the bottom to the top was the brown ion. **(c, right)** Cartoon representation of the NavSp1 structure showing one frame during the simulation indicated by an arrow on the left panel, with three ions bound within the permeation pathway shown as spheres. **(d, left)** MD simulation trajectories showing chloride ion binding events and binding occupancy at the coiled coil residue R236. Each individual chloride ion that occupies this location is shown in a different color. **(d, right)** Cartoon representation of the NavSp1 structural showing one frame during the simulation indicated by an arrow on the left panel, with a chloride ion bound to R236 shown as a sphere. **(e, left)** The sum of the interface predicted template modeling (ipTM) and the predicted template modeling (pTM), which measure the accuracy of the predicted inter-subunit model interfaces and the predicted model, respectively, was similar in all models with pores in open or closed conformations. **(e, right)** The 5$^{th}$-9$^{th}$ ranked models with a closed pore did not exhibit drastically different conformations of the CTD, as shown by the opposing chain alpha carbon distances plotted for all residues in the C-terminus for these top models.

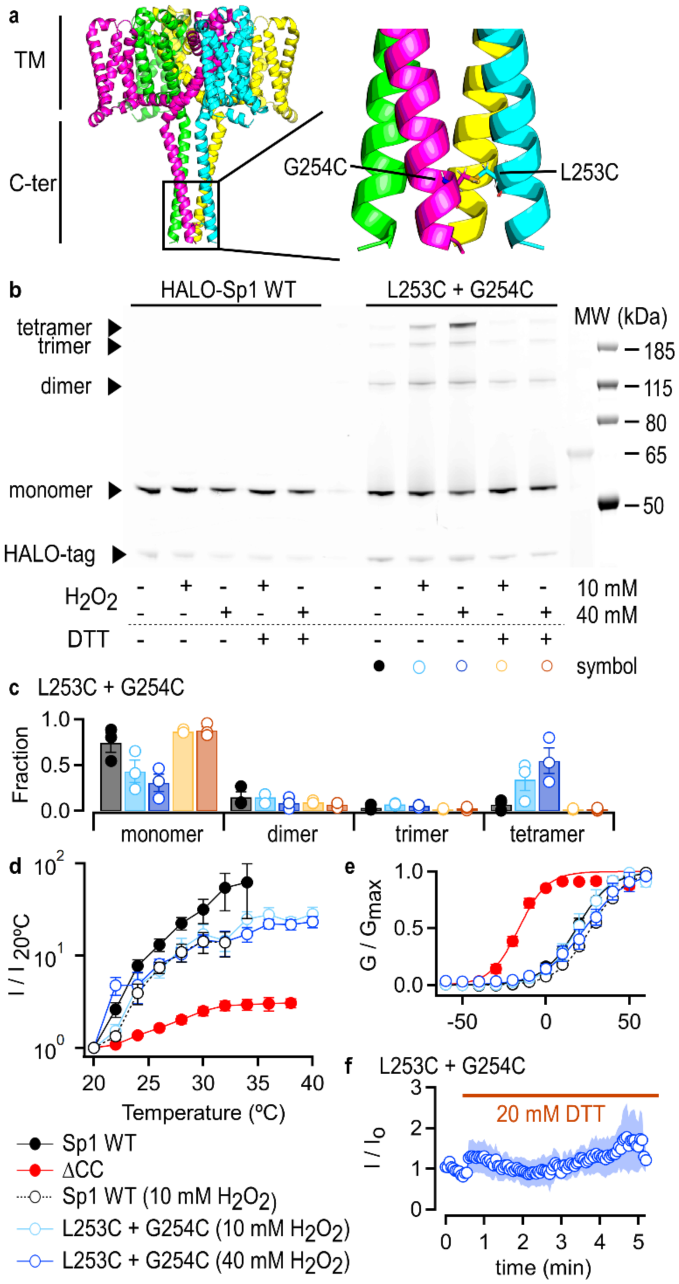


**Figure S4. The integrity of the coiled coil is preserved in the response of the Na$_V$Sp1 channel to heat. (a)** AlphaFold2 model of NavSp1 L253C+G254C. **(b)** In-gel JF549 dye fluorescence after non-reducing SDS-PAGE of lysates from cells expressing Halo-tagged NavSp1 channel constructs. Experimental conditions before lysis for each lane are denoted at the bottom. Molecular weights (in kDa): Halo-tag (35.6), monomer (66.4), dimer (132.8), trimer (199.2), and tetramer

(265.7). **(c)** In-gel fractions of Halo-NavSp1 L253C+G254C complexes from individual experiments (circles) as in (b) and their mean ± SEM (bars, n = 3). **(d)** Current-temperature relations (mean ± SEM; WT NavSp1 + $H_2O_2$, n = 4; HALO-NavSp1 L253C+G254C, 10 mM $H_2O_2$, n = 6; 40 mM, n = 5). **(e)** Conductance-voltage relations at room temperature (mean ± SEM; WT NavSp1 + $H_2O_2$, n = 6; Halo-NavSp1 L253C+G254C, 10 mM $H_2O_2$, n = 4; 40 mM, n = 4). Continuous curves in (c) and (e) are fits to the Boltzmann equation. **(f)** Time-course of Halo-NavSp1 L253C+G254C-mediated currents before and after exposure to DTT. Cells were pre-treated with 40 mM $H_2O_2$. Currents were elicited by voltage steps from -90 to 30 mV at room temperature. Data shown as mean ± SEM (n = 5).

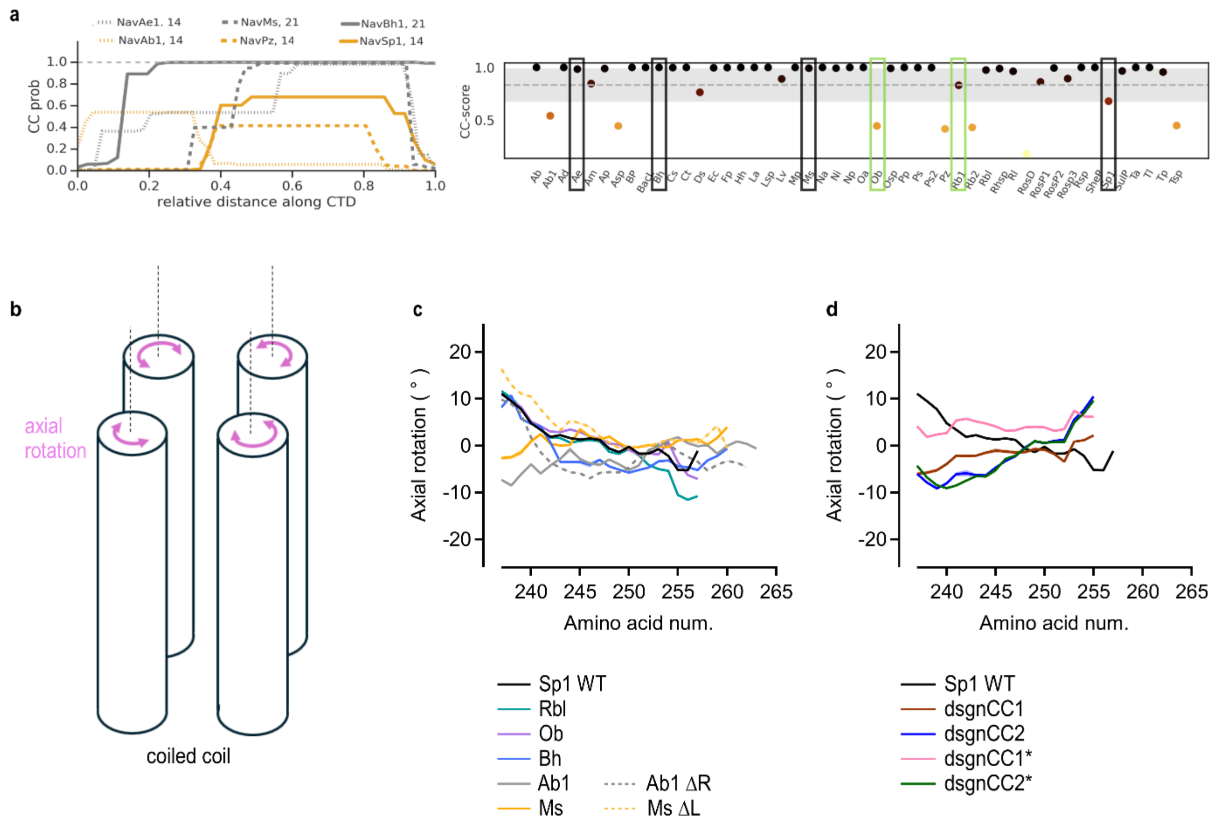


**Figure S5. Coiled coil properties. (a)** Plots of the coiled coil probability (49) (CC prob) of the full CTD for distinct the BacNavs used in this study to generate coiled coil chimeras with the NavSp1 proximal CTD and transmembrane domain. The calculated CC probability for NavSp1 (continuous yellow curve) was lower than that of NaChBac (NavBh1), NavMs, and NavAe1. Before we carried out experiments for this study, we had initially predicted that this observation could be related to the temperature sensitivity of each of the channels, where channels with strong temperature dependence would have lower CC probabilities than channels where temperature is not sufficient to disrupt CTD integrity. Thus, Ob and Rbl (highlighted in green boxes on the right) were chosen because they had different coiled coil probability prediction scores (right panel), which we calculated as the maximal value of any CC probability for any window according to the algorithm from Lupas et al (49). **(b)** Cartoon representing a homotetrameric parallel coiled coil as in NavSp1, showing the structural significance of the axial rotation parameter as calculated by samCC (34). **(c, d)** Mean axial rotation per amino acid calculated by samCC (34) for each of the channel chimeras or WT Sp1, whose identity is denoted by the colors.

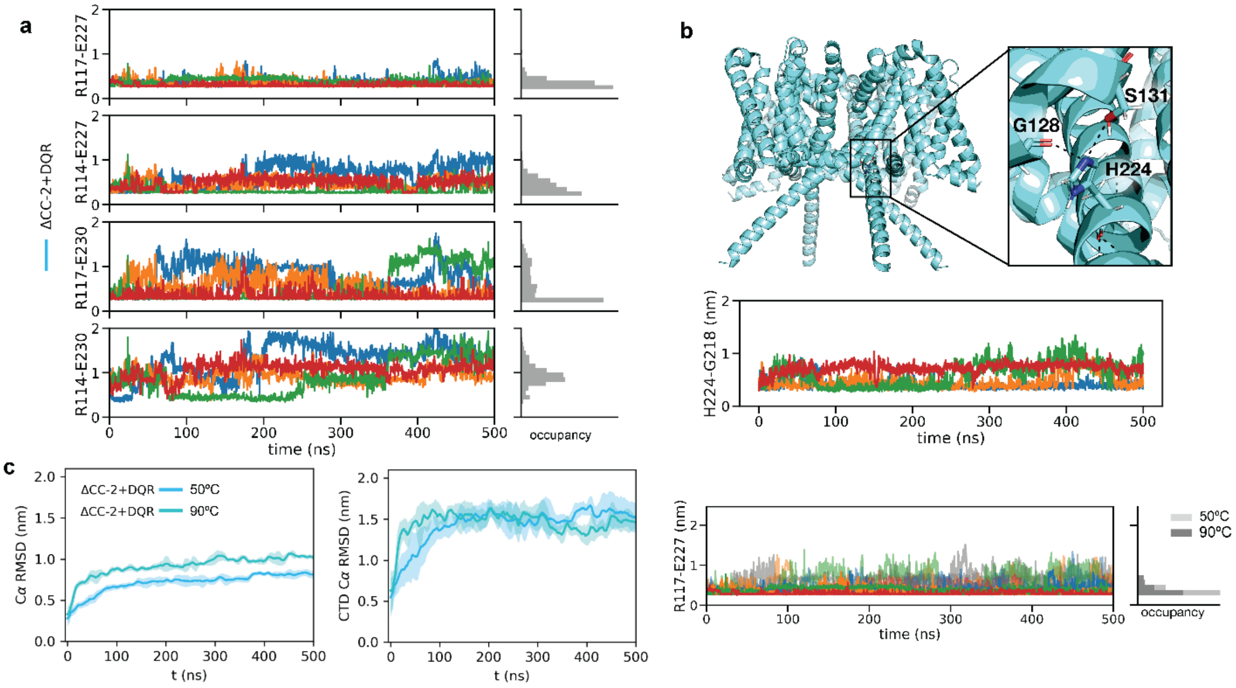


**Figure S6. Analysis of interactions between the C-terminus and the transmembrane domain of Na$_V$Sp1. (a)** Closest heavy atom distance in nm between residues in the proximal C-terminus (E227 and E230) and the S4-S5 linker (R117 and R114) shown for a single MD simulation started from the open state of the ΔCC-2 +DQR construct with a splayed truncated coiled coil, colored by chain. To the right, a histogram of all three replicates over 500ns of each of these distances is shown, mean and standard deviation for each distance across all three trajectories (in nm): R117-E227: mean = 0.3529 with std = 0.11, R114-E227: mean = 0.4468 with std = 0.20, R117-E230: mean = 0.5854 with std = 0.36, and R114-E230: mean = 0.9582 with std = 0.28. **(b, top)** Top-ranked open pore model for the ΔCC-2+DQR (cyan) with a splayed C-terminus, showing additional interactions between the proximal C-terminus and the S4-S5 linker. **(b, bottom)** Closest heavy atom distance between residues H224 and G128 during MD simulations started from the open state of the ΔCC-2 +DQR construct. **(c, left)** Windowed-averages of RMSD of all alpha-carbons for the ΔCC-2 +DQR construct comparing simulations performed at 50 ºC and 90 ºC, showing that the 90 ºC simulations indeed deviate more from the starting structure but remain stable. **(c, middle)** Windowed-averages of RMSD of alpha-carbons for only the C-terminal domain show that the 50 ºC and 90 ºC simulations have similar behavior within the CTD region, though the 90 ºC simulations deviate faster from the starting conformation. **(c, right)** The closest-heavy atom distance between residues R117 and E227 shows that the higher temperature representative trajectory (shown here in grey) does not deviate substantially from the lower temperature simulation (shown colored by chain for a representative simulation) – indicating that his interaction can remain stable at high temperatures. To the right, histograms of all replicates are shown. For the lower temperature, the mean = 0.3529 nm and std = 0.1105 nm. For the higher temperature, mean = 0.4104 nm and std = 0.1772 nm.

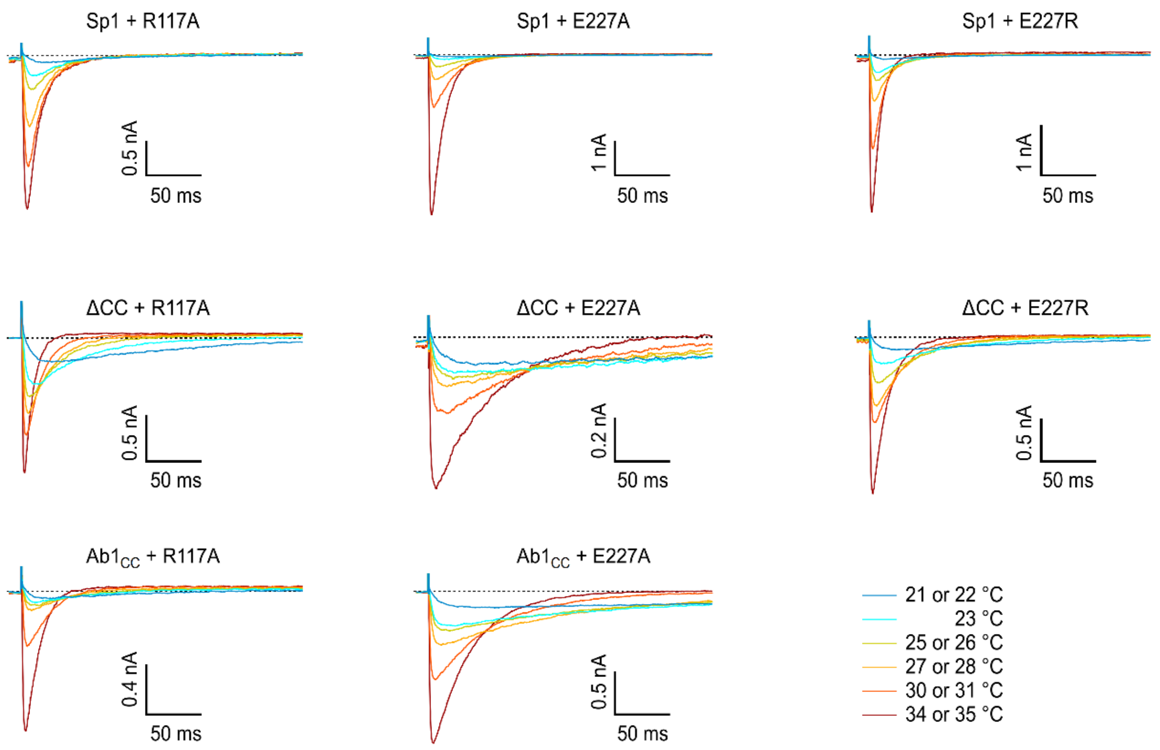


**Figure S7. Experimental testing of interactions between the C-terminus and the transmembrane domain of Na$_V$Sp1.** Representative current traces elicited during a temperature ramp by repetitive voltage steps from -90 to 30 mV in cells expressing Na$_V$Sp1 channel constructs. Current traces for only six temperatures are shown for visual clarity.

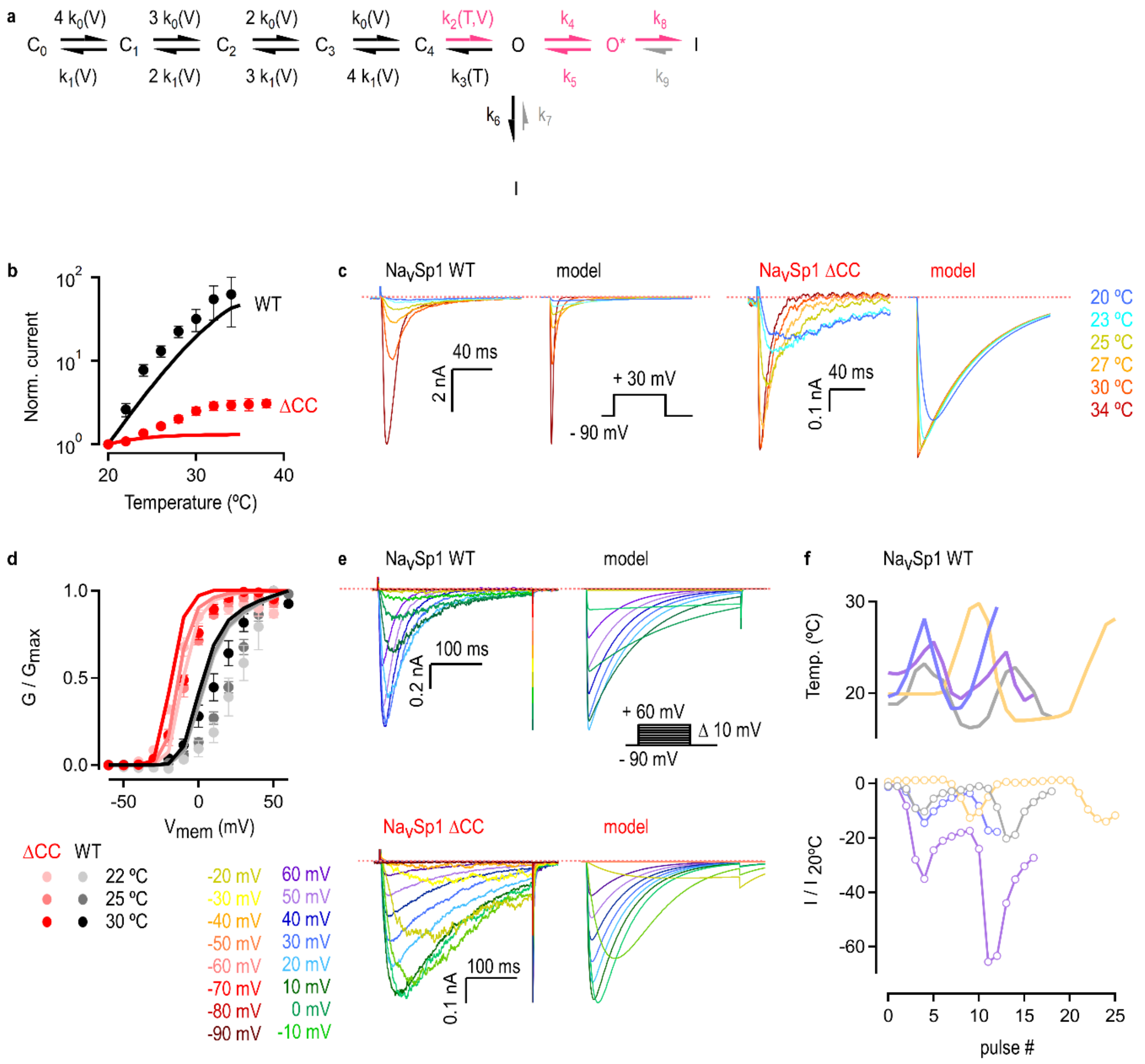


**Figure S8. Mathematical model and reversibility of the temperature-dependent activity of $Na_VSp1$ channels. (a)** Scheme of the model used to describe channel activity. C are closed states, O are open states, and I is the inactivated state. Forward and reverse rate constants and their dependencies on voltage or temperature are shown. Recovery from inactivation (grey symbols) did not contribute to our calculations. States and transitions colored in magenta are those that are influenced by the CTD. The parameters used to model currents are: WT, $k_0(0) = 4{,}000$ s$^{-1}$; $k_1(0) =$ $1{,}000$ s$^{-1}$; $k_2 = 15{,}000$ s$^{-1}$; $k_3 = 1{,}000$ s$^{-1}$; $k_4(0) = 800$ s$^{-1}$; $k_5(0) = 0.00001$ s$^{-1}$; $z_0 = 1.5$; $z_1 = -1.5$; $z_2$ $= 0.1$; $\Delta H_2 = 250{,}000$ kJmol$^{-1}$; $\Delta S_2 = 0.790$ kJmol$^{-1}$K$^{-1}$; $\Delta H_3 = 5{,}000$ kJmol$^{-1}$; $\Delta S_3 = 1$ kJmol$^{-1}$K$^{-1}$; $\Delta$CC, $k_0(0) = 4{,}000$ s$^{-1}$; $k_1(0) = 1{,}000$ s$^{-1}$; $k_2(0) = 250{,}000$ s$^{-1}$; $k_3 = 1{,}000$ s$^{-1}$; $k_4(0) = 800$ s$^{-1}$; $k_5(0)$ $= 0.00001$ s$^{-1}$; $k_6 = 50{,}000$ s$^{-1}$; $k_7 = 10$ s$^{-1}$; $k_8 = 15$ s$^{-1}$; $k_9 = 0.00001$ s$^{-1}$; $z_0 = 1.5$; $z_1 = -1.5$; $z_2 = 0.1$;

$\Delta H_2$ = 250,000 kJmol$^{-1}$; $\Delta S_2$ = 0.790 kJmol$^{-1}$K$^{-1}$; $\Delta H_3$ = 5,000 kJmol$^{-1}$; $\Delta S_3$ = 1 kJmol$^{-1}$K$^{-1}$; $V_{rev}$ = 75 mV. **(b)** Experimental (symbols) and calculated (continuous curves) current-temperature relations from experiments as in (c), for WT (black) or ΔCC (red) NavSp1 channels. **(c)** Comparison between experimental and calculated current traces elicited by voltage steps from -90 to 30 mV during a temperature ramp as in Fig.1g for WT or ΔCC NavSp1. The dotted red line denotes the zero-current level. Current traces for only six temperatures are shown for visual clarity. **(d)** Comparison between calculated (continuous curves) and experimentally determined (symbols) conductance-temperature relations obtained from data as in (e) for WT or ΔCC NavSp1. **(e)** Comparison between experimental and calculated current families at room temperature elicited by a series of voltage steps between -90 and +60 mV. **(f)** Peak WT NavSp1 current magnitude as a function of time and in response to the changes in temperature depicted on the top panel. Colors denote independent experiments. Currents were elicited by voltage steps from -90 to 30 mV.
